# Cell Morphology accurately predicts the nuclear shape of adherent cells

**DOI:** 10.1101/2024.12.28.630588

**Authors:** Sebastian Lawton, Rosaline A. Danzman, Renzo Spagnuolo, Skylar Stephan, Sydney Graul, O’Neil Wiggan, Soham Ghosh, Ashok Prasad

**Affiliations:** School of Biomedical Engineering; Cellular and Molecular Biology Program; Department of Chemical and Biological Engineering; Department of Biochemistry and Molecular Biology; Department of Mechanical Engineering Colorado State University, Fort Collins, Colorado 80523 USA

## Abstract

Cells are internally tensed, or prestressed, largely by actomyosin contractility. We hypothesized that nuclear shape is quantitatively predictable from cell shape since prestress couples them both. We trained machine learning models on a publicly available image database of the WTC-11 cell line and predicted shape modes of the nucleus with high accuracy. We develop a U-Net architecture-based model, Cell2Nuc, that predicted nuclear voxels from the cell membrane with accuracies between 74%-87%. To investigate prestress, we cultured and imaged HeLa cells after inhibiting actomyosin contractility. The Cell2Nuc model retrained on the HeLa cells predicted nuclear voxels with slightly lower accuracy. Statistical analysis revealed changes in nuclear size and chromatin organization upon prestress inhibition. Similar trends were seen in images taken from NIH3T3 cells. Thus, cell shape encodes features of nuclear shape, their coupling is partly due to actomyosin contractility, whose abrogation leads to changes in chromatin organization of mechanosensitive origin.

## Introduction

The shape of the nucleus plays a critical role in regulating cellular function by influencing genome organization, gene expression, and mechanotransduction ^1–3^. Nuclear deformations are important in normal physiological processes ^2^, such as in hematopoietic stem and progenitor cells (HSPCs) differentiation into the myeloid lineage ^4^ or in cell fate determination in the embryo ^5^. Aberrations in nuclear shape are critical signatures of disease, such as in laminopathies ^6^ or cancer ^7^. In both diseased and healthy cell, deformations of the nucleus have been shown to causally correlate with changes in gene expression ^2,8,9^.

Nuclear deformations are mechanical in origin and arise through the transmission of forces between the cytoskeleton and the nuclear envelope that it is strongly coupled with ^9–13^. As the primary force generating system in most cells due to the action of non-muscle myosin II, the actin cytoskeleton is perhaps the most important cellular component involved in nuclear shaping^12^, though other cytoskeletal constituents also play a role ^14^, Actomyosin contractility can deform the nucleus and has been shown to be associated with nuclear dysmorphia in cancer ^15–18^. Actomyosin networks pull, rotate and deform the nucleus in migrating cells ^19–21^. When cells are geometrically constrained, actomyosin networks shape both the cell and the nucleus^11,22,23^. Actomyosin networks are also key determinants of cell shape ^24^, and thus couple cell shape to nuclear shape. This coupling between cell and nuclear shape through mechanics and the cytoskeleton suggests that nuclear shape may be quantitatively predictable from cell shape. This would imply that cell shape also encodes nuclear shape, which may be part of the reason why cell shape is so informative about cell state, as has been demonstrated previously by several groups ^25,26^.

How could forces shape the nucleus unless the cell was moving or dividing? The cytoskeleton exists in a state of prestress largely due to the action of myosin II motors on the actin cytoskeleton ^27^. How the state of prestress impacts the nucleus is less well understood. If we assume, like much prior work on nuclear mechanics, an elastic model of the nucleus, the resting ground state of the nucleus is approximately spherical and deformations from this shape require application of forces or steric constraints ^28,29^. In an alternative model, the nuclear drop model, forces shape the nucleus but are not required for its maintenance ^30^. The former model implies a strong coupling between cell and nuclear shapes while the latter would predict a weaker coupling in non-motile cells.

To our knowledge this is the only study that tries to explicitly predict the shape of the nucleus from the shape of the cell at a single cell level. That there is an approximately constant ratio between the size of cells and the size of the nucleus (called the karyoplasmic ratio) has been known for a long time ^31^. However, this tells us nothing about nuclear shape and positioning. The co-determination of nuclear shape and cell shape was studied using very different methods in Ref. ^32^ who constructed a “predictive model of nuclear shape given cell shape (or vice versa) using the weighted average nuclear shapes of neighbors in cell shape space” ^32^. Viana et. al. ^33^ who took some of the images we analyze here carried out some correlation analysis of their data, showing that geometric descriptors of shape such as surface area and volume of the cell and nucleus were highly correlated, reaching R squared values in the range of 0.73 to 0.90. In this work we systematically study the coupling between cell shape and nuclear shape by utilizing deep artificial and convolutional neural networks that have been designed to predict nuclear shape from cell shape alone, as well as classical statistical models. We build our initial models by training them on a large, publicly available dataset of cell and nuclear images taken by Viana et. al. ^33^ from the Allen Institute of Cell Sciences. To understand the role of cytoskeletal prestress, specifically actomyosin contractility, we culture Hela cells that have been transformed by labeling myosin II with GFP, and image these cells after treatment with the ROCK inhibitor Y-27632, the ROCK inhibitor H-1152 and the myosin II inhibitor, blebbistatin. We retrain our machine learning on this dataset and carry out a systematic statistical analysis of two-dimensional parameters. To validate some of our observations on another cell line, we also cultured and imaged NIH3T3 cells under the same experimental perturbations.

## Results

### Nuclear shape parameters can be predicted by cell shape parameters in WTC-11 cell lines

The Allen Institute has published a database ^33^ of over two hundred thousand images of the hiPSC WTC-11 cell line (STAR Methods). Using the metadata, we picked the interphase cell sample dataset with cell and nuclear masks available. The dataset contained images of several intracellular components but here we use only the images of the cell membrane and the nuclear envelope. Apart from the single cell images, the dataset also reported the decomposition of cell membrane and nuclear envelope images into a spherical harmonic expansion (SHE), performed separately for the cell and nucleus. To reduce the dimensionality of the SHE we performed a principal component analysis (PCA) on each set separately.

We chose the first 20 principal components to describe the cell membrane shapes (covering 86.3% of total variation) and the first 12 principal components for the nuclear shape (covering 91.5% of total variation). These principal components are denoted shape modes in this paper, and they reduced the dimensionality of the dataset by an order of magnitude. The percent variance contributed by each of these shape modes can be seen in Table S1 (Supplementary Information).

We then asked whether key features of nuclear shape were predictable from cell shape modes using regression models. We trained an artificial neural network (ANN) regression model to predict nuclear height, nuclear area and nuclear volume, as well as each of the 12 nuclear shape modes from all 20 of the cellular shape modes. We used 10-fold cross validation for testing the accuracy of the trained ANN model. The average correlation across all the 10-test folds between the predicted nuclear parameter and the actual nuclear parameter are shown in Table 1, and scatter plots are shown in Fig. S1 in the Supplementary Information. As the column labeled “Shape Modes 1-20” shows, nuclear height was predicted very well with a correlation of 0.938. Nuclear area and volume were also predicted well with correlations of 0.879 and 0.917 respectively, and so were the first five shape modes, with correlations above 0.7. The remaining shape modes were predicted with correlations between 0.5 to 0.7. Overall, the model could explain about 65% of the total variance in the data.

**Table 1:**
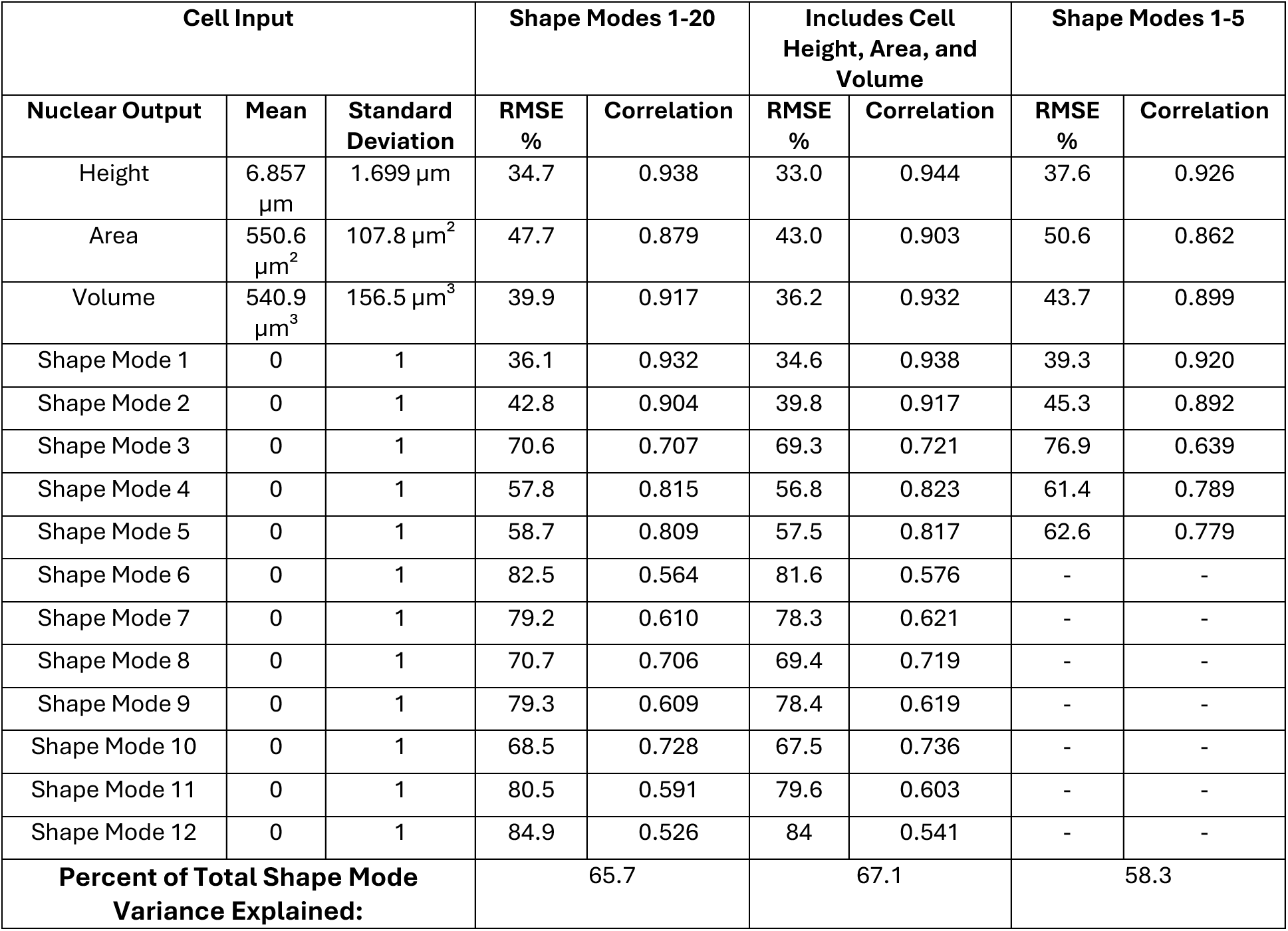
Performance Comparison of ANN models on hiPSCs. The average root-mean squared error (RMSE) as a percentage of the standard deviation and correlations for each of their predicted nuclear output variables is shown. The R² value for each model on the shape modes is also reported as the percent of the total shape mode variance explained.

We wondered if the regression models would do better if they were additionally supplied with cell height, volume and area as explanatory variables. Indeed, this ANN model performed marginally better (Table 1, column labeled “Includes Cell Height, Area and Volume”) in predicting the nuclear shape parameters. Finally, we wondered whether all 23 explanatory variables were required, and we trained an ANN regression model with only 5 cell shape modes as explanatory variables (Table 1, column “Shape Modes 1-5”). To our surprise, correlations between the predicted and actual values remained high for the truncated model, for some variables almost as high as for the model with all 20 shape modes. However, the percentage of variance explained dropped to 58%.

We next asked whether a nonlinear model was critical for the prediction accuracy. Three multiple linear regression (MLR) models were created alongside the neural networks using the same explanatory variables as a baseline comparison for model performance (Table 2). We found that prediction accuracy as measured by percentage of variance explained dropped by roughly 5% for the larger models, but only by 2% for the truncated model. Correlation coefficients dropped only a little for every dependent variable in the MLR models. Thus, nonlinearity contributes to model accuracy but is not critical, since linear models perform well.

**Table 2:** Performance of MLR models on hiPSCs. The average root-mean squared error (RMSE) as a percentage of the standard deviation and correlations for each of their predicted nuclear output variables is shown. The R² value for each model on the shape modes is also reported as the percent of the total shape mode variance explained.

| Cell Input |  |  | Shape Modes 1-20 |  | Includes Cell Height, Area, and Volume |  | Shape Modes 1-5 |  |
| --- | --- | --- | --- | --- | --- | --- | --- | --- |
| Nuclear Output | Mean | Standard Deviation | RMSE % | Correlation | RMSE % | Correlation | RMSE % | Correlation |
| Height | 6.857 $\mu\text{m}$ | 1.699 $\mu\text{m}$ | 44.1 | 0.897 | 33.4 | 0.942 | 45.3 | 0.891 |
| Area | 550.6 $\mu\text{m}^2$ | 107.8 $\mu\text{m}^2$ | 59.7 | 0.802 | 43.6 | 0.900 | 61.3 | 0.791 |
| Volume | 540.9 $\mu\text{m}^3$ | 156.5 $\mu\text{m}^3$ | 52.0 | 0.854 | 36.9 | 0.930 | 53.2 | 0.847 |
| Shape Mode 1 | 0 | 1 | 42.4 | 0.906 | 35.5 | 0.935 | 43.0 | 0.903 |
| Shape Mode 2 | 0 | 1 | 47.4 | 0.881 | 41.1 | 0.909 | 47.5 | 0.880 |
| Shape Mode 3 | 0 | 1 | 78.4 | 0.620 | 76.9 | 0.639 | 79.1 | 0.612 |
| Shape Mode 4 | 0 | 1 | 59.8 | 0.802 | 61.4 | 0.789 | 62.8 | 0.778 |
| Shape Mode 5 | 0 | 1 | 62.6 | 0.780 | 62.6 | 0.780 | 64.0 | 0.769 |
| Shape Mode 6 | 0 | 1 | 90.5 | 0.424 | 90.5 | 0.424 | - | - |
| Shape Mode 7 | 0 | 1 | 84.3 | 0.539 | 84.6 | 0.539 | - | - |
| Shape Mode 8 | 0 | 1 | 74.3 | 0.669 | 75.3 | 0.700 | - | - |
| Shape Mode 9 | 0 | 1 | 84.3 | 0.533 | 84.6 | 0.533 | - | - |
| Shape Mode 10 | 0 | 1 | 76.3 | 0.646 | 69.9 | 0.716 | - | - |
| Shape Mode 11 | 0 | 1 | 84.6 | 0.533 | 82.5 | 0.565 | - | - |
| Shape Mode 12 | 0 | 1 | 89.9 | 0.438 | 89.9 | 0.438 | - | - |
| **Percent of Total Shape Mode Variance Explained:** |  |  | 63.4 |  | 60.2 |  | 56.2 |  |

To visually assess how good the model predictions were, we used the predicted nuclear shape modes to reconstruct the images. The average of the predictions made by the 10 cross-fold models was used in the reconstruction. Figure 2 displays the true nucleus image, the reconstruction from the true SHE which represents ground truth for the ML models, and the reconstructed images from the predicted shape modes from the ANN and the MLR models for one sample cell. The figure shows that the SHE representation lacks fine detail of nuclear surface due to truncation of the terms, though the major trends of the nuclear shape are captured, such as being tilted, squashed, or stretched. For example, the nucleus is a little flattened on one end in the XZ plane, but the SHE reconstruction does not capture this, but it does accurately capture the tilt. ANN and MLR predictions look quite similar and generally capture these major features in the SHE representation.

**Figure 1.**
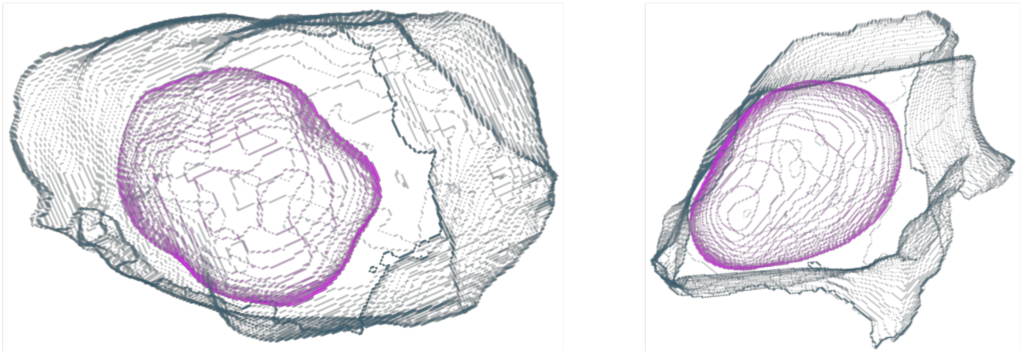
Representative examples of cells and nuclei from the Allen Institute database. The cell membrane is shown in gray and the nuclei in pink. The cell membrane image was used to predict the nucleus image.

**Figure 2.**
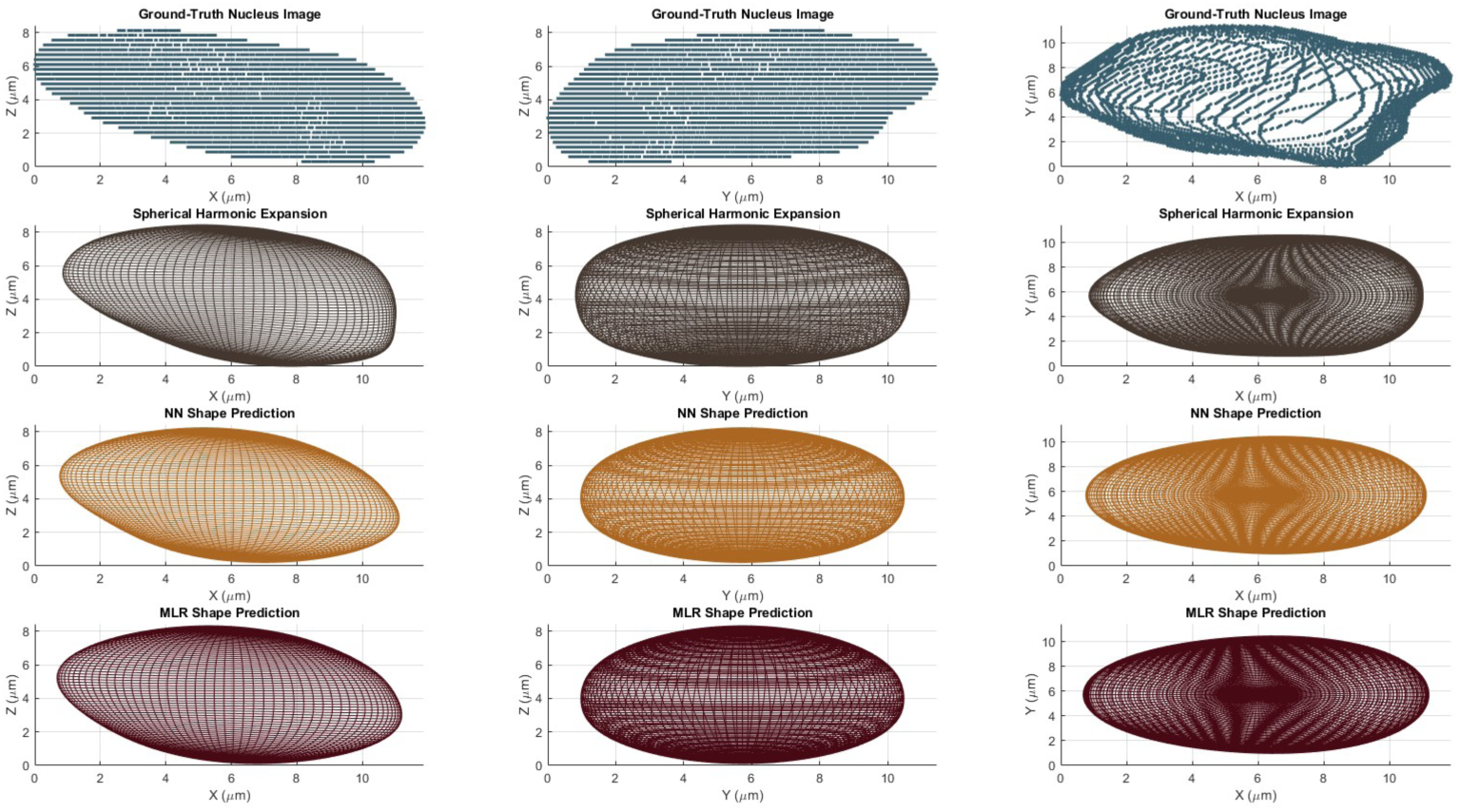
Nuclei images, reconstructions and predictions in different planes. Top row: XZ, YZ and XY projections of the actual nucleus. Second row: SHE reconstruction using truncated expansion. Third row: Reconstructed image from ANN models. Fourth row: reconstructions from MLR models.

### A Vox2Vox style U-Net model can reconstruct nuclear shape from cell shape

We next asked if we could predict nuclear shapes directly from the 3D shape data instead of a SHE representation, using a convolutional neural net. We chose a U-Net architecture for this purpose since it is a standard architecture for cell segmentation ^34^, for example in the Vox2Vox brain image segmentation model of Cirillo et. al. ^35^, and can work with smaller datasets. However, we applied the U-Net architecture in a novel way, as we explain below.

A standard U-Net architecture follows an encoder-decoder structure, resembling a “U” shape, with symmetrical contracting and expanding paths. The encoder progressively captures contextual information by applying convolutional layers followed by max-pooling operations, reducing the spatial dimensions while increasing feature depth. In typical applications, the decoder then reconstructs the segmentation map by applying upsampling operations, concatenating high-resolution features from the encoder (via skip connections), and further convolutions ^34^. While we preserved this basic idea, in our case the decoder arm maps the features learnt by the encoder from the cell membrane images to predict the voxels that form the nucleus. In other words, instead of mapping each voxel to a class label, the model identifies a different set of voxels predicted to be occupied by the nucleus based on cell membrane shape. We thereby describe it as the Cell image to Nuclear image (Cell2Nuc) model (Supplementary Figure S2).

A subset of 3,824 cells with segmented cell and nuclear mask images in interphase from the Allen database were utilized for Cell2Nuc model training and prediction. The Dice-Sørensen coefficient or dice score that is a ratio of correctly predicted voxels over total voxels was used to assess the accuracy of predictions and the models. We also calculate the RMSE and the R^2^ value derived from it. Ten-fold cross validation averages of dice scores as well as other accuracy metrics are reported in Table 3.

**Table 3:** Accuracy of Cell2Nuc model on hiPSC dataset. The Dice score is a ratio of correctly predicted voxels over total voxels and lies between 0 and 1. The translated image has been shifted to coincide with the true nucleus center as described in the text. See STAR Methods for all formulae and methods of calculation.

| Measure | Mean | Standard Deviation | $R^2$ | RMSE % |
| --- | --- | --- | --- | --- |
| Dice score of raw prediction | 0.735 | 0.058 | - | - |
| Dice score of binary prediction | 0.806 | 0.073 | - | - |
| Dice score of translated image | 0.865 | 0.061 | - | - |
| Translated distance (as % of height) | 16.60 | 12.10 | - | - |
| Height ($\mu\text{m}$) | 7.875 | 1.707 | 0.735 | 51.6 |
| Surface Area ($\mu\text{m}^2$) | 520.4 | 98.2 | 0.224 | 92.5 |
| Volume ($\mu\text{m}^3$) | 553.4 | 154.0 | 0.344 | 83.9 |
| Spherical Parameter S | 0.177 | 0.059 | 0.844 | 39.4 |
| Flattening Parameter F | 0.779 | 0.150 | 0.803 | 44.4 |
| Angle of Tilt $\theta$ ($^\circ$) | 23.00 | 12.92 | -0.337 | 115.6 |

The initial U-Net results with a high threshold for prediction gave us a dice score of 73.5% which indicated that the model was accurately identifying the majority of nuclear voxels. Adjusting this threshold increased the dice score to 80.6% (Dice score of binary prediction). Visual inspection of the predicted nuclear images showed that the model was consistently making errors in estimating the nuclear center. We therefore translated the predicted image to the real nuclear center and calculated the dice score, which then was even higher, at 86.5% (Dice score of translated image). Note that in medical imaging a dice score of 70% is treated as a good score ^36,37^, so our scores are quite significant by those criteria. An animation of a prediction near the mean Dice score of the translated binary images is shown in Figure 3, and additional animations of good and bad predictions are shown in Supplementary Figure S5 and S6. A large variety of animations showing examples of good and bad predictions based on shape measures and dice scores can be found in the image repository (see Data Availability).

**Figure 3.**
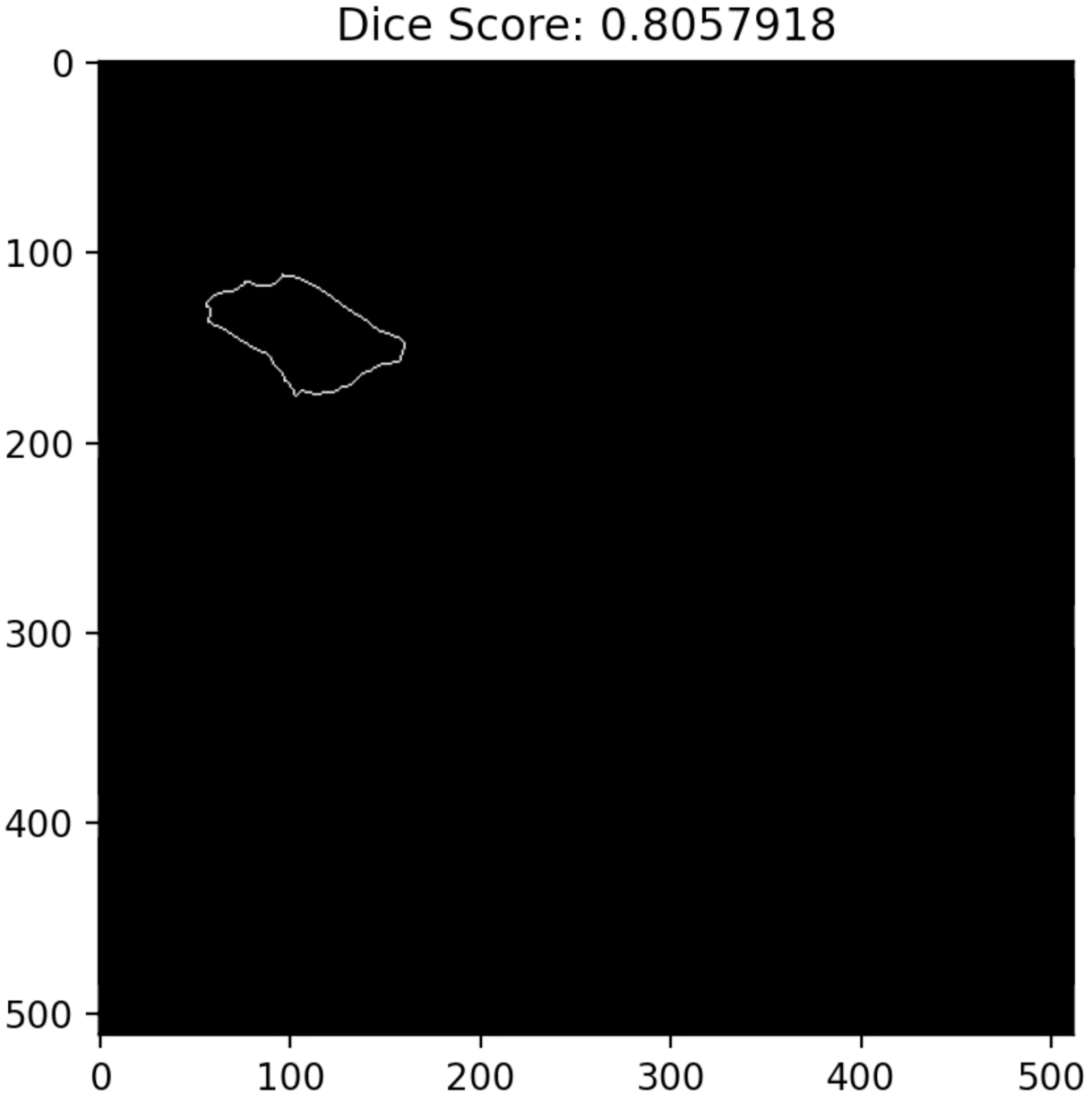
An animation of a prediction produced by the Cell2Nuc model near the mean Dice score of the binary images. The animation shows the z-stacks of the image starting from the top. The cell membrane is the white boundary, and the nucleus, predicted and actual, are the solid interior pixels. The predicted nucleus has been translated so that the center of the ground truth nucleus matches its own. White pixels represent the ground truth nucleus that were not predicted as such by the model (false negatives). Magenta pixels pertain to the overlap of the ground truth and predicted nuclei (true positives). Green pixels pertain to the predicted nucleus which does not overlap the ground truth image (false positives).

The height of these nuclei was predicted with a high R² similar to that of the ANN models, though the surface area and volume dimensions were not. This type of prediction task is more difficult than that of the ANN, especially when we consider that the U-Net models had a smaller training set (of sample size of 3824 rather than 197,682 for the ANNs) and that these models were not directly trained on the true values of these dimensions.

We then asked if the reconstruction accurately captured second-order features of nuclear shape, such as its sphericity, flattening and tilt. The spherical parameter S measures how far an object differs from a sphere based upon its height, surface area, and volume measures. We saw that S was quite low, indicating that the nuclei were generally close to a sphere in shape, and that the models were able to achieve a high R² with their predictions. Similarly, the flattening parameter F showed that these nuclei were generally flatter than a perfect sphere, and the model was able to reach a high R² with this parameter too.

The angle of tilt Θ was not well predicted by these models. A negative R² indicates that the predictions performed worse than a model which only predicts the mean for each sample, increasing the RMSE % to greater than the standard deviation. Here, it may be the case that the calculation of this angle was too naïve an estimation and that a more sophisticated measure could achieve better results, as the high Dice scores would imply that the models tend to follow trends like the movement of nuclei cross sections through the z stacks.

Visual inspection of several predictions indicated that the Cell2Nuc model was predicting more irregular shapes than the SHE-based models, and thus it could be picking up surface features of the nucleus better than the former. We next asked if there were other, second-order shape features that the ANN models were doing better at.

### ANN models can predict non-trivial features of nuclear shape

We reasoned that each individual shape mode must be capturing some non-trivial features of nuclear shape, as indeed shown for other types of shape modes ^33,38^. So, we plotted each of the nuclear shape modes to further investigate what aspect of the nuclear shape they account for in the dataset.

A 3D mesh was created representing the extreme ends (z score = ±2) of each shape mode, with each other mode held constant at its mean value (z score = 0). In Figure 4 the average cell and nuclear mesh is displayed in three profiles as a reference. Here we can directly see that the average nuclear shape does indeed appear to be generally spherical in shape with a squished or flatter height, confirming the measurements taken in the 3D images. Figure 5 shows the cell shape mode extremes and Figure 6 shows those of the nucleus in their most informative profile, as assessed by visual inspection.

**Figure 4.**
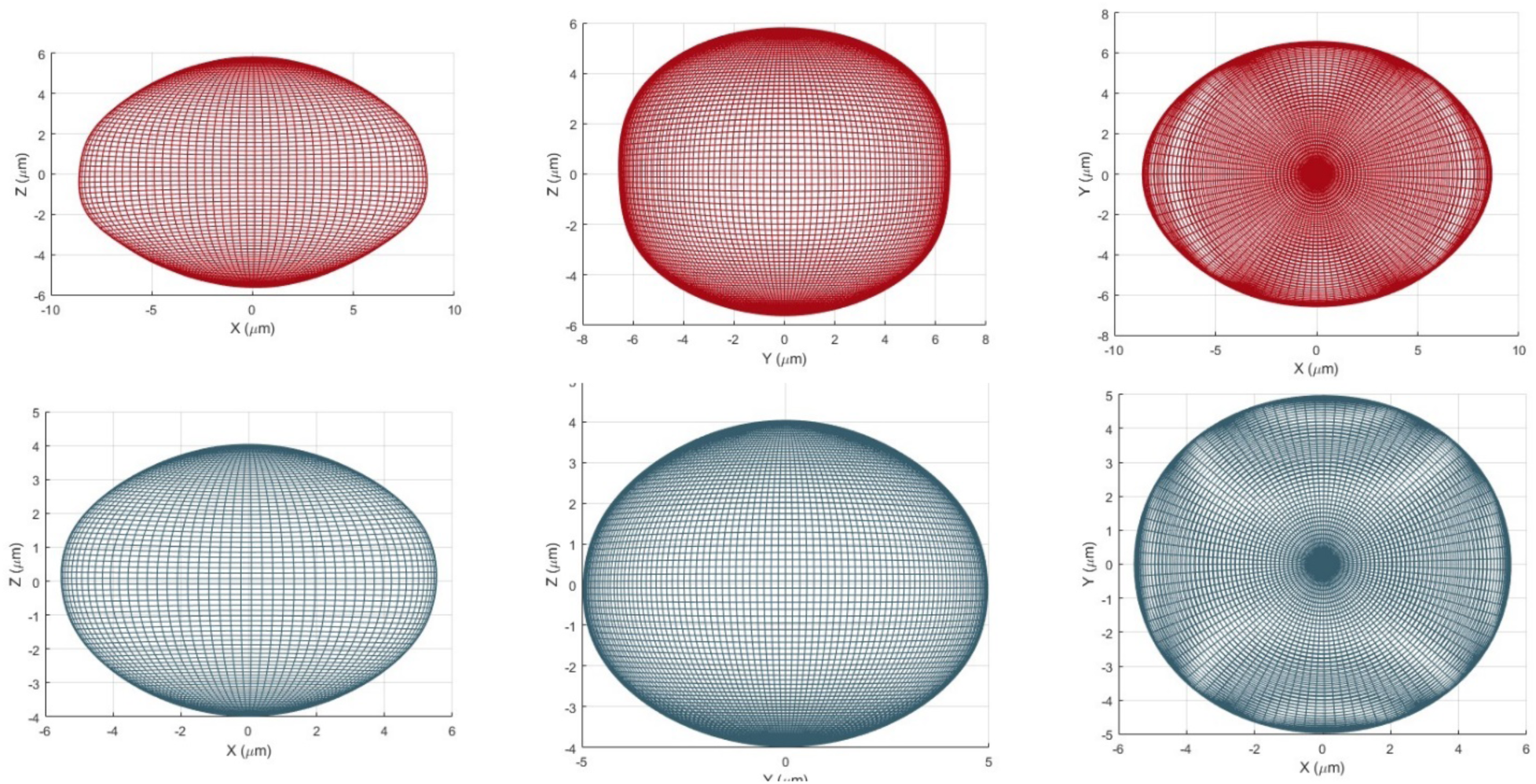
Average cell (red, above) and average nucleus (blue, below). 3D meshes were reconstructed using the mean shape modes and are shown here in three profiles, XZ, YZ and XY from left to right.

**Figure 5.**
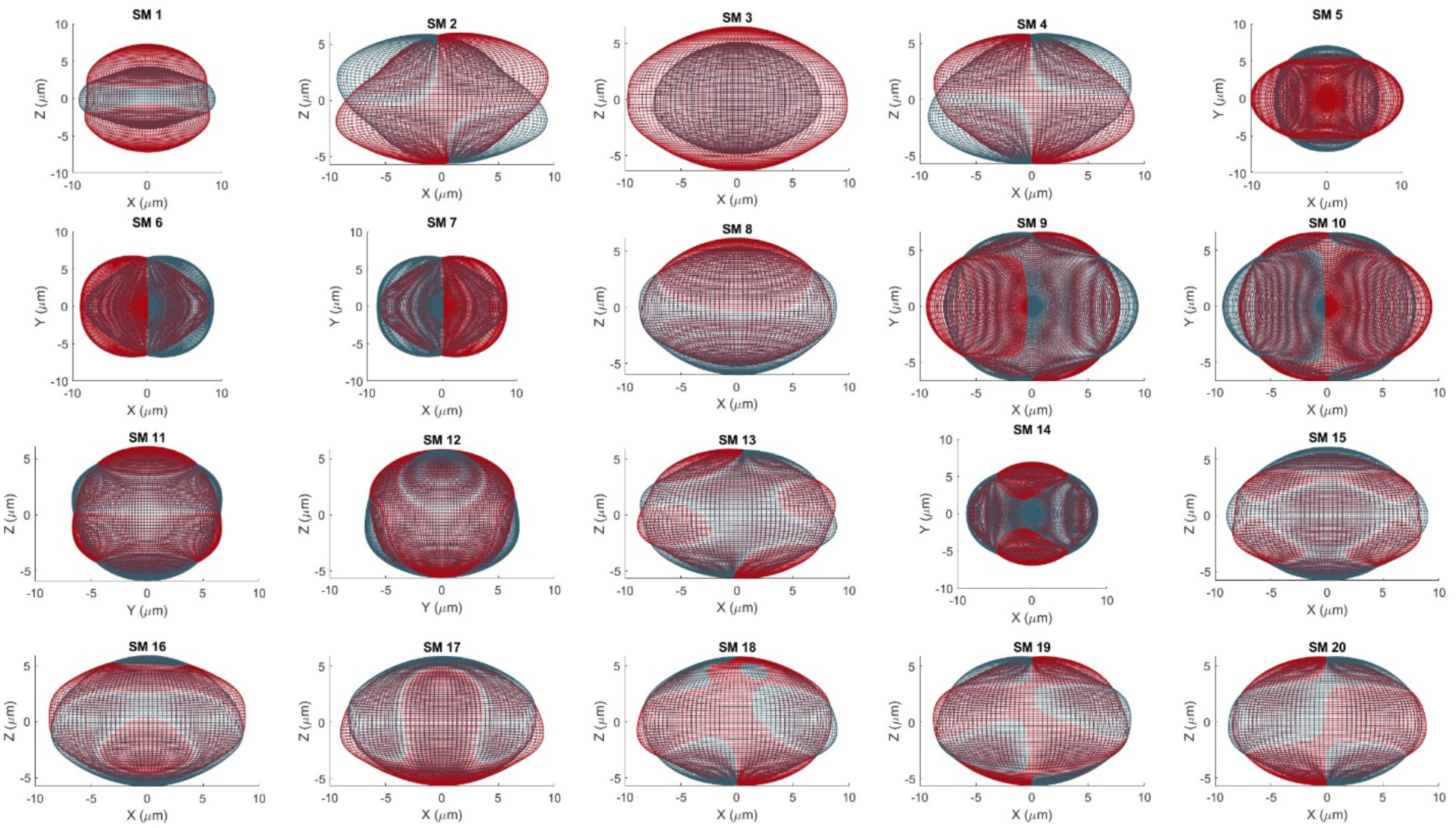
Cell Shape modes visualized by reconstructing the cellular image using z= +2 or -2 for each shape mode, while holding all others at the mean of 0. These positive (red) and negative (blue) extremes of the 20 cell shape modes are shown in their most informative 3D profile.

**Figure 6.**
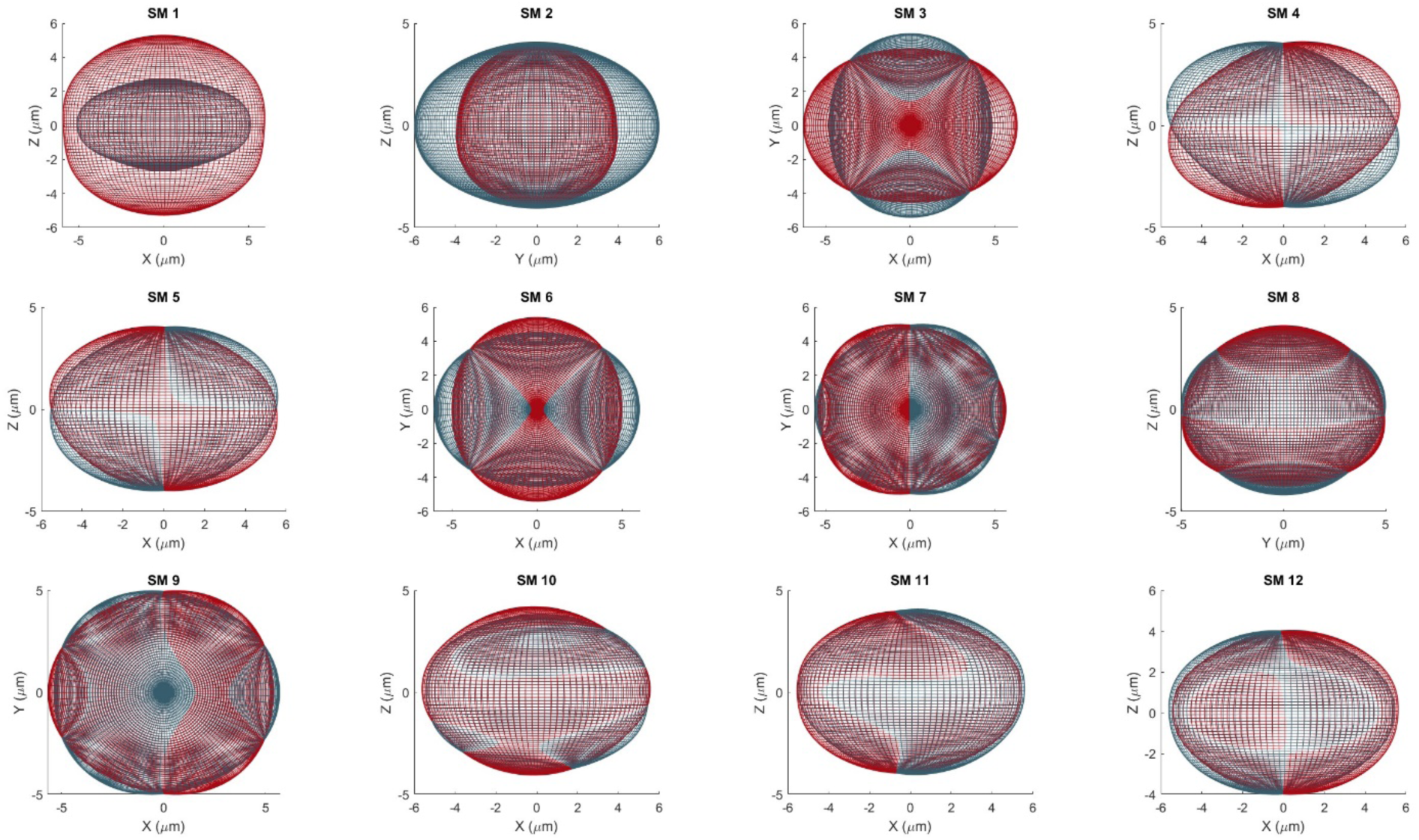
Nuclear shape modes visualized by reconstructing the cellular image using z= +2 (blue) or z= -2 (red) for each shape mode while holding others constant. The images were examined in all three cartesian planes and are plotted here in their most informative 3D profile.

Figure 6 allows us to interpret some of the shape modes qualitatively. Nuclear Shape Mode 1 (NSM1) describes the overall volume of the nucleus and captures the variation in height, as seen by the projections in the XZ and YZ planes shown in Figure 7A. Since the MLR model performance is similar to the ANN models, we can use the decomposition of total variance (see STAR Methods) in the MLR models to assess which cell parameters are most important in predicting each specific nuclear shape mode (Supplementary Table S2). For NSM1 it turns out that the top three cell shape modes (CSM) are CSM1, CSM3 and CSM5. All three of these seem to capture different dimensions of overall size (Fig. 5).

**Figure 7.**
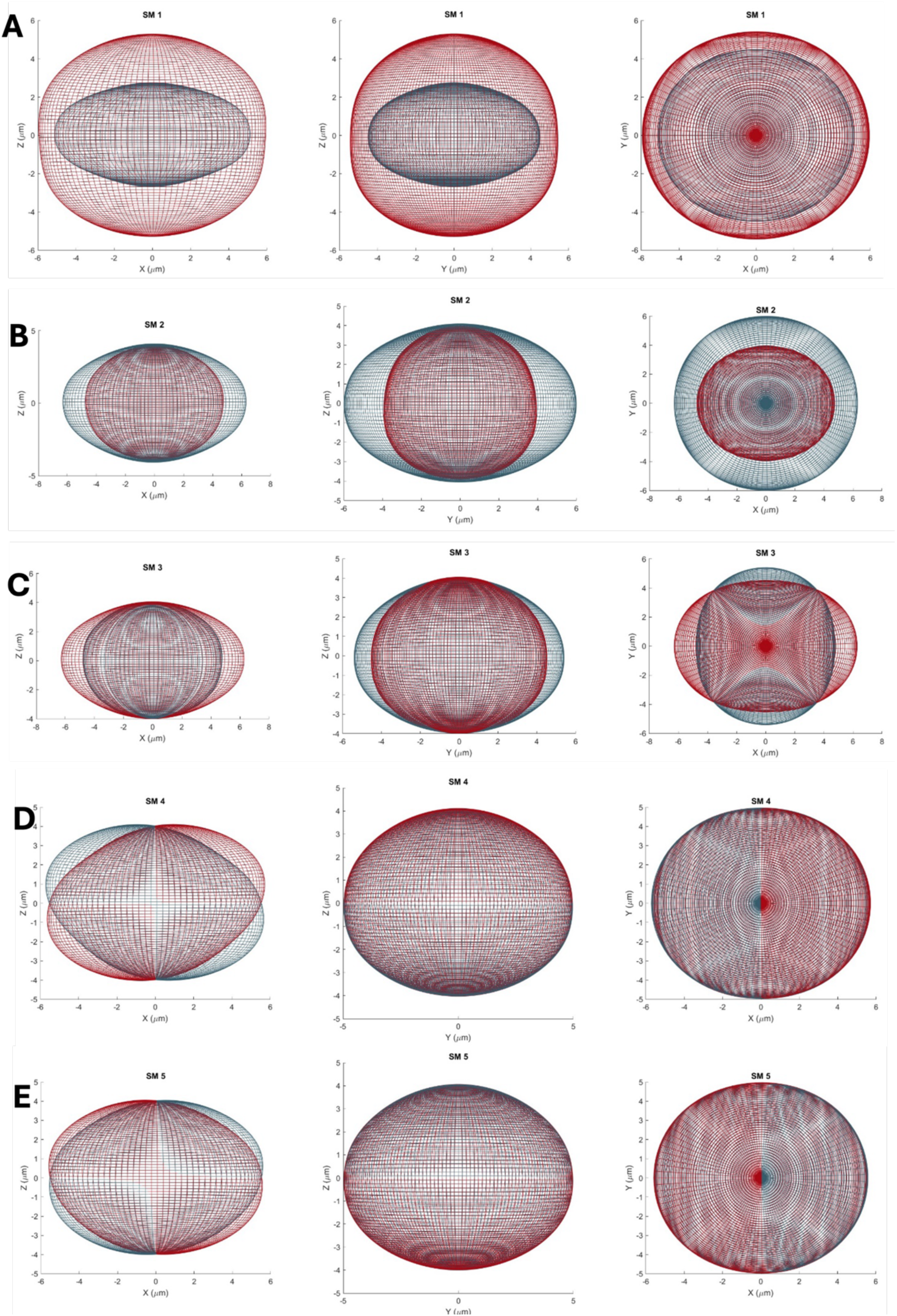
Projections of the first 5 shape modes (A-E) in the XZ (first column), YZ (second column) and XY (third column) planes.

NSM2, displayed in Figure 6 and Figure 7B, captures changes in the cross-sectional area of the nucleus in the XY plane, which is not captured by NSM1. Here again, the primary factors for prediction were CSM1, CSM3 and CSM5 along with the volume and height dimensional data (Supplementary Table S2). NSM3, shown in Figure 7C, describes the major and minor axis lengths and orientation of the nucleus in the XY-plane. This shape mode was a little harder to predict accurately compared with all of the first 5 shape modes, with a correlation coefficient of 0.707 and an RMSE of about 70% (Table 1). It is not clear why this shape mode should have a lower accuracy. The main contributing parameter is CSM5 (Supplementary table S2).

The last two of the first 5 nuclear shape modes, NSM4 NSM5, in Figures 7D and 7E respectively, appear to be complementary in describing the tilt of the nucleus in the XZ plane. It should be noted that these nucleus images were aligned with their major axis in the XY plane lying along the x-axis in order to calculate the SHE coefficients in a uniform way by Viana et. al. ^33^. It is particularly interesting to see that this phenomenon of nuclear tilt appears to occur only in the XZ plane, i.e. only in the plane of the major axis. The primary predictor of NSM4 was CSM2, and secondly CSM4, and the opposite was true for NSM5.

For NSM6 and higher modes, there is a substantial drop off in the accuracy of the ANN predictions. Interestingly, NSM6 appears complementary to NSM3, but its best predictors were CSM12 and CSM14. The higher shape modes of either the nucleus or the cell are harder to interpret qualitatively.

In summary, cell shape encodes several nontrivial features of nuclear shape, which can be predicted with reasonably high accuracy. We next asked whether cell and nuclear shapes are mechanically linked together through the prestress in the actin cytoskeleton. To answer this question, we cultured HeLa cells expressing GFP-labeled myosin II and imaged them in 3D using a laser scanning confocal microscope, after staining actin and chromatin as detailed in the STAR Methods. We then segmented the cell and nuclear images in 3D using CellPose ^39^ and CellProfiler ^40^ and analyzed the images using the Cell2Nuc U-Net model as well as standard statistical methods. We then treated the HeLa cells with the ROCK inhibitor Y-27632, the ROCK inhibitor H-1152 and the myosin II inhibitor, blebbistatin. We again tested the Cell2Nuc U-Net models on the cells treated with the cytoskeletal drugs.

### The Cell2Nuc U-Net model predicts over 70% of nuclear voxels correctly for HeLa cells

The Cell2Nuc models trained on the Allen Institute cells did not perform well when applied directly to the HeLa cell images, but when retrained after starting from previously trained models, they converged rapidly, despite the smaller size of the dataset (between 1,072 and 2,148 samples per treatment). However, prediction accuracy on the HeLa cells was generally worse than on the hiPSCs (Table 4) and the trends were similar. The average dice score for the initial prediction was 67.5% over the 10 test sets in 10-fold cross-validation, and this improved to 75% after adjusting the threshold and 80% after translating the nuclear center. However, despite predicting 75-80% of the pixels correctly, the models could not estimate the variations in nuclear parameters accurately, as seen by the large RMSE compared with the standard deviation. Some reasons for this could be a smaller dataset and lower image resolution as compared with the Allen Institute dataset.

**Table 4.** Accuracy of the Cell2Nuc U-Net model on HeLa cells. Numbers reported are averages of 10-fold cross-validation.

| Group | Measure | Mean | Standard Deviation | RMSE % |
| --- | --- | --- | --- | --- |
| Control Group | Dice score of raw prediction | 0.675 | 0.107 | - |
|  | Dice score of binary image | 0.750 | 0.124 | - |
|  | Dice score of translated image | 0.803 | 0.093 | - |
|  | Translated distance (as % of height) | 20.56 | 16.87 | - |
| | Height ($\mu\text{m}$) | 6.782 | 1.696 | 99.0 |
| | Surface Area ($\mu\text{m}^2$) | 430.7 | 82.8 | 95.3 |
| | Volume ($\mu\text{m}^3$) | 478.2 | 135.4 | 94.7 |
|  | Spherical Parameter S | 0.195 | 0.064 | 100.7 |
|  | Flattening Parameter F | 0.701 | 0.144 | 121.6 |
| | Angle of Tilt $\Theta$ ($^\circ$) | 6.816 | 7.665 | 108.3 |

Though prediction accuracy dropped for all treated cells, as can be seen in Table 5, it is noteworthy that the dice score for the binary images and the translated images remained higher than 70% even in these cases. There was a general decrease in prediction accuracy for shape features which remained higher than the standard deviation in almost all cases. The one exception was the H-1152 treated cells. For H-1152, the nuclear height and the S and F parameters were able to achieve higher R² and lower RMSE values.

**Table 5.** Accuracy of the Cell2Nuc U-Net models for HeLa cells treated with cytoskeletal drugs.

| Group | Measure | Mean | Standard Deviation | RMSE % |
| --- | --- | --- | --- | --- |
| Blebbistatin | Dice score of raw prediction | 0.666 | 0.107 | - |
|  | Dice score of binary image | 0.730 | 0.143 | - |
|  | Dice score of translated image | 0.786 | 0.124 | - |
|  | Translated distance (as % of height) | 21.15 | 15.82 | - |
| | Height ($\mu\text{m}$) | 6.078 | 1.044 | 123.9 |
| | Surface Area ($\mu\text{m}^2$) | 380.5 | 62.0 | 140.7 |
| | Volume ($\mu\text{m}^3$) | 393.8 | 96.7 | 131.9 |
|  | Spherical Parameter S | 0.204 | 0.052 | 118.6 |
|  | Flattening Parameter F | 0.673 | 0.104 | 132.6 |
| | Angle of Tilt $\Theta$ ($^\circ$) | 6.155 | 6.691 | 112.2 |
| Group | Measure | Mean | Standard Deviation | RMSE % |
| H-1152 | Dice score of raw prediction | 0.662 | 0.109 | - |
|  | Dice score of binary image | 0.734 | 0.136 | - |
|  | Dice score of translated image | 0.800 | 0.105 | - |
|  | Translated distance (as % of height) | 20.47 | 18.39 | - |
| | Height ($\mu\text{m}$) | 6.787 | 1.537 | 88.7 |
| | Surface Area ($\mu\text{m}^2$) | 382.3 | 72.6 | 107.4 |
| | Volume ($\mu\text{m}^3$) | 395.6 | 113.4 | 104.2 |
|  | Spherical Parameter S | 0.176 | 0.062 | 84.2 |
|  | Flattening Parameter F | 0.752 | 0.156 | 98.4 |
| | Angle of Tilt $\Theta$ ($^\circ$) | 5.456 | 5.651 | 116.0 |
| Group | Measure | Mean | Standard Deviation | RMSE % |
| Y-27632 | Dice score of raw prediction | 0.627 | 0.106 | - |
|  | Dice score of binary image | 0.685 | 0.152 | - |
|  | Dice score of translated image | 0.759 | 0.135 | - |
|  | Translated distance (as % of height) | 25.89 | 18.50 | - |
| | Height ($\mu\text{m}$) | 6.005 | 1.278 | 101.1 |
| | Surface Area ($\mu\text{m}^2$) | 374.4 | 61.9 | 152.5 |
| | Volume ($\mu\text{m}^3$) | 385.5 | 97.1 | 143.6 |
|  | Spherical Parameter S | 0.209 | 0.060 | 112.0 |
|  | Flattening Parameter F | 0.668 | 0.122 | 127.3 |
| | Angle of Tilt $\Theta$ ($^\circ$) | 6.875 | 7.606 | 112.0 |

Application of the U-Net model to HeLa cells suggested that cell and nuclear shapes continued to be coupled together, and we were unable to see significant differences between the control and treatment groups. However, this could be because the prediction accuracy was insufficient. Analysis of the 3D images also showed that the treatment groups had a smaller height and nuclear volume when compared with the control group (Tables 4 and 5). This observation led us to ask whether signatures of contractility disruption could be seen in maximum projection two-dimensional images too.

### Actomyosin contractility disruption results in changes of nuclear size but not shape in 2D

We analyzed maximum projection fluorescent images of HeLa cells (STAR Methods) using parameters calculated by CellProfiler as well as custom python scripts. Disruption of actomyosin contractility with Blebbistatin (inhibiting non-muscle myosin II), Y-27632, or H-1152 (both Rho Kinase inhibitors) in HeLa cells, led to a small decrease in all 2D nuclear size parameters. Specifically, nuclear area, perimeter, minor and major axes decreased in response to disrupted actomyosin contractility. Interestingly, there was little to no observed change in other shape parameters such as roundness (aspect ratio), circularity and eccentricity. While nuclear size changed there was almost no change in cell size or shape parameters in response to the drug treatment (Supplementary Figure S3).

Distribution of nuclear area across the four treatment groups shows a higher mean for the control group at 203 µm compared to each of the groups with impaired contractility (Figure 8). The decrease in nuclear area was most pronounced in the H-1152 group with a mean of 183µm. All differences are significant with p-values between 1.19 ×10^−22^ to 1.11×10^−11^.

**Figure 8.**
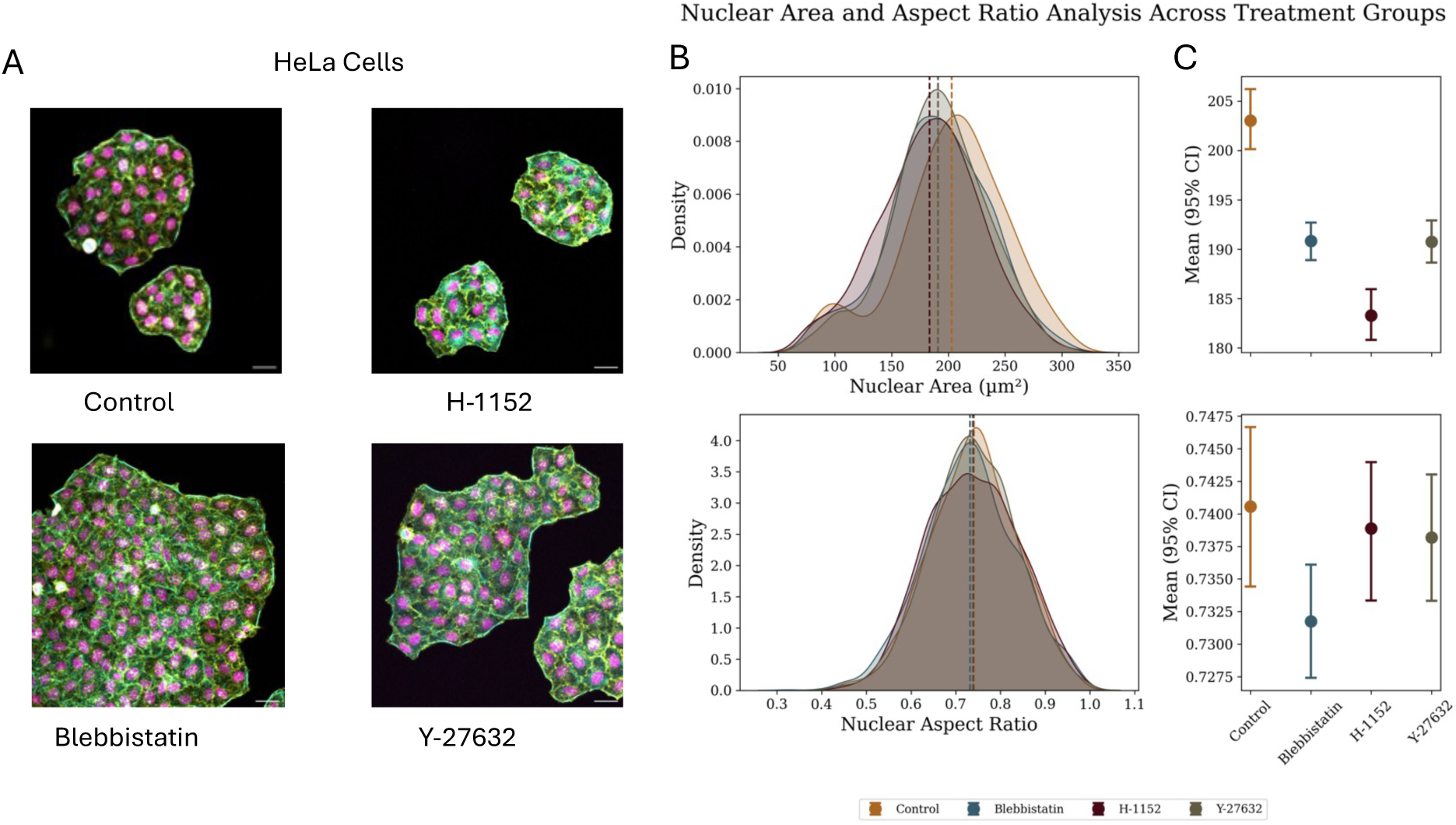
Representative HeLa cell images for each treatment group and select nuclear feature distributions and means with 95% CI. A. Fluorescence microscopy images of HeLa cells expressing myosin-II-GFP (Yellow) stained with DAPI (Magenta) and phalloidin (Cyan) for each treatment group. Scale bars = 20µm. B. Density distribution of nuclear area (top) and aspect ratio (bottom) by treatment group. C. Mean nuclear area (top) and mean aspect ratio (bottom) with 95% CI by treatment group.

While nuclear size (area, perimeter, major and minor axis) also decreases with disruption of actomyosin contractility (Figure 8 and Supplementary Figure S4), the overall shape of the nucleus remains largely unaffected, as illustrated by nuclear aspect ratio shown in Figure 8. The mean aspect ratio for all four experimental groups (0.73-0.74) are very close, and the differences are not significant.

Our protocol involves fixing the cells after a 30 minute incubation of the contractility disrupting drug. Thus, the changes observed are likely mechanical in origin. Our results are consistent with the hypothesis that cytoskeletal prestress actively maintains nuclear size, but they also imply that prestress acts in a largely isotropic manner.

Disruption of cytoskeletal prestress could also disrupt the mechanical coupling between the cell and nucleus. Though we were unable to find direct evidence of this using the Cell2Nuc U-Net model, we tested this idea using regression models on 2D data.

### Cell shape parameters can predict nuclear dimensions in 2D

We first carried out simple linear regressions between individual cell and nuclear morphological features extracted from 2D images of HeLa cells. Broadly speaking measures of cell size had significant linear correlations with measures of nuclear size, with R^2^ values for many feature pairs at around 0.59 (Supplementary Table S3). Inhibition of actomyosin contractility with Blebbistatin, Y-27632, and H-1152 lowered many of these correlations for feature pairs with R^2^ values just above about 0.4. The one exception to that trend is the effect of H-1152 that in some cases increased the correlation between cell and nucleus feature pairs (Supplementary Table S3).

We then performed a multiple linear regression (MLR) analysis using ridge regression, with 18 cell features as well as their squares and products to capture nonlinearities as explanatory variables. The regression R^2^ for the seven nuclear features of equivalent diameter (the diameter of a circle having the equivalent area of the nucleus measured), median radius, mean radius, nuclear area, nuclear perimeter, nuclear minor axis and nuclear major axis were between 0.5 and 0.7 for the control cells, but fell significantly for all of the treatment cells (Figure 9C). The decline was most marked for blebbistatin treated cells. As before, shape features such as eccentricity, solidity and compactness were poorly predicted (Figure 9C). These results thus indicate that cell size and nuclear size is strongly coupled in control cells and this coupling is decreased when contractility is disrupted.

**Figure 9.**
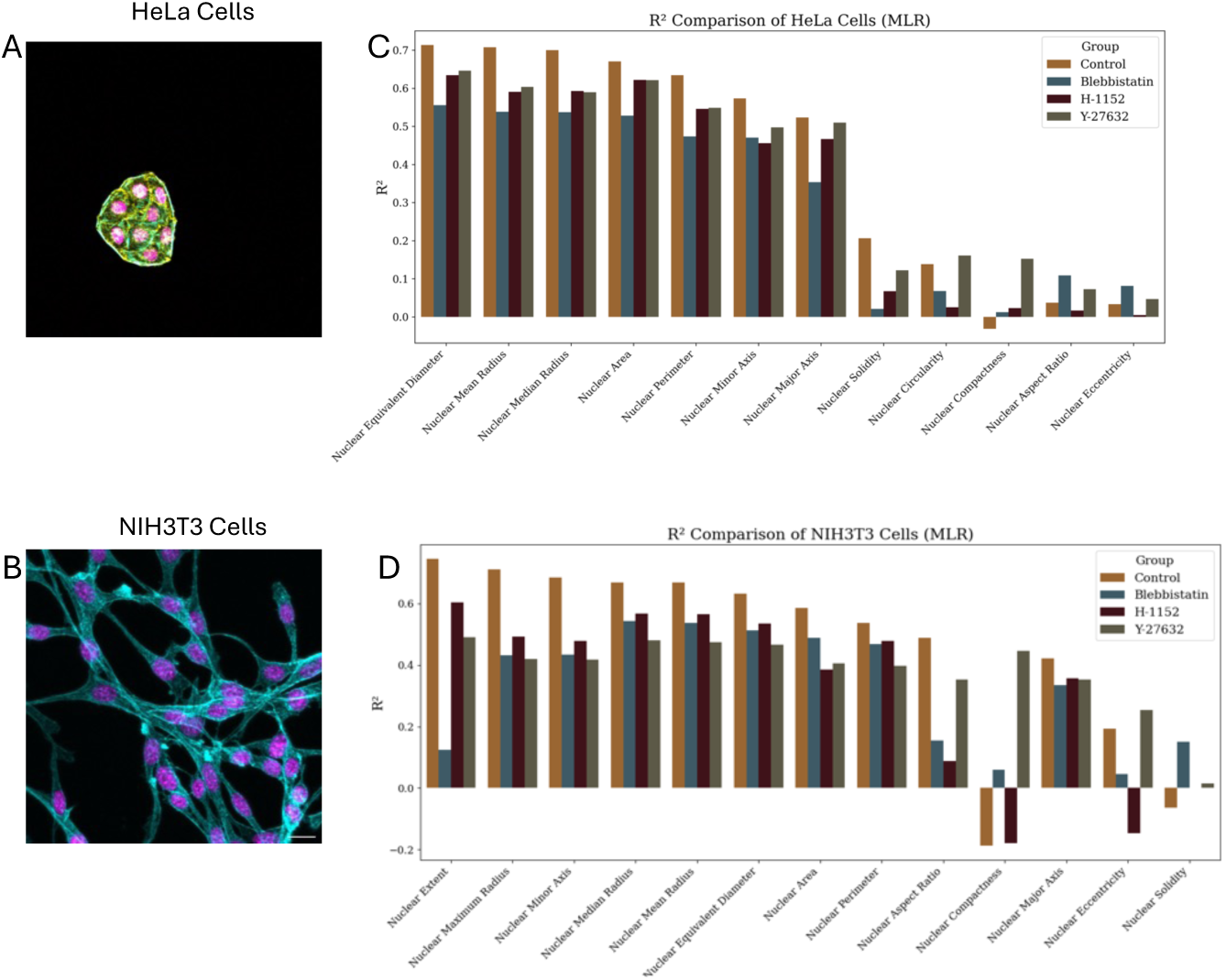
Representative fluorescence microscopy images with Multiple Linear Regression (MLR) results predicting nuclear features from cell features. Representative images of HeLa (A) and NIH3T3 cells (B). Scale bar = 20µm. R^2^ values predicting a subset of nuclear features from cell features for HeLa (C) and NIH3T3 cells(D) by treatment group.

We were unable to explain nuclear shape features like compactness, solidity and eccentricity from cell shape parameters. Previous groups have found that elongated cells typically had elongated nuclei ^41^. However, the HeLa cells in our culture were generally quite rounded or polygonal, and perhaps therefore nuclear elongation was uncorrelated with cell elongation. We speculated that cells with very different morphology may not display the same cell-nucleus relationships seen in HeLa cells. To test this idea, we cultured and imaged NIH3T3 fibroblasts (experimental details in STAR Methods).

### Similar patterns of prediction are reflected in cells with vastly different morphology

While adherent HeLa cells have been described as “egg-like” alluding to their rounded shape ^42^, NIH3T3 fibroblasts exhibit vastly different morphology with spindle or stellate shapes (Figure 9A and B). We cultured NIH3T3 fibroblasts and imaged them as we did HeLa cells. Despite their very different morphology, the results of the Multiple Linear Regression analysis of cell shape – nuclear shape coupling were very similar to the HeLa cells. Thus, nuclear size parameters were predictable from cell shape parameters but nuclear shape parameters such as elongation were not well predicted. The MLR models have higher predictive power for control cells, and the predictability dropped significantly when actomyosin contractility was disrupted (Figure 9D). The highest R^2^ values for the control group was 0.71 for nuclear maximum radius, which dropped to 0.49 for H-1152 treated cells, 0.43 for Blebbistatin, and 0.42 for Y-27632 (Figure 9D). As before, the more complex shape-based features such as solidity were poorly predicted.

### Disruption of actomyosin contractility leads to chromatin redistribution in a cell-line specific manner

Previous groups have shown that changes in nuclear shape and size may impact gene expression ^43^ and this is accompanied by chromatin reorganization. We asked whether there was evidence of chromatin reorganization even in the short time frames of <30 minutes after drug treatment. To test this, we calculated the Chromatin Condensation Parameter (CCP) ^44^ that detects the most obviously condensed chromatin using Sobel edge detection (Figure 10). The CCP has lower values when the chromatin is more diffuse and higher values for more condensed chromatin. We found that in HeLa cells the CCP increased after disruption of actomyosin contractility in all of the treatment groups (Figure 10A). The CCP was the lowest in control cells, and it was the highest in Blebbistatin treated cells. Conover’s non-parametric test shows significant difference between the control group compared with all other groups (p-values between 0.0085 to 8.94×10^17^ ) as well as a significant difference between the blebbistatin group compared to the other two treatments for HeLa cells (p-values 7.83 ×10^8^ for H-1152 and 2.24×10^14^ for Y-27632, Supplementary Table S4). NIH3T3 fibroblasts on the other hand showed a decrease in chromatin compaction when cells were treated with H-1152 and an increase when cells were treated with Y-27632, with no change for blebbistatin-treated cells (Figure 10B). Thus abrogation of actomyosin contractility leads to changes in nuclear size as well as changes in chromatin organization.

**Figure 10.**
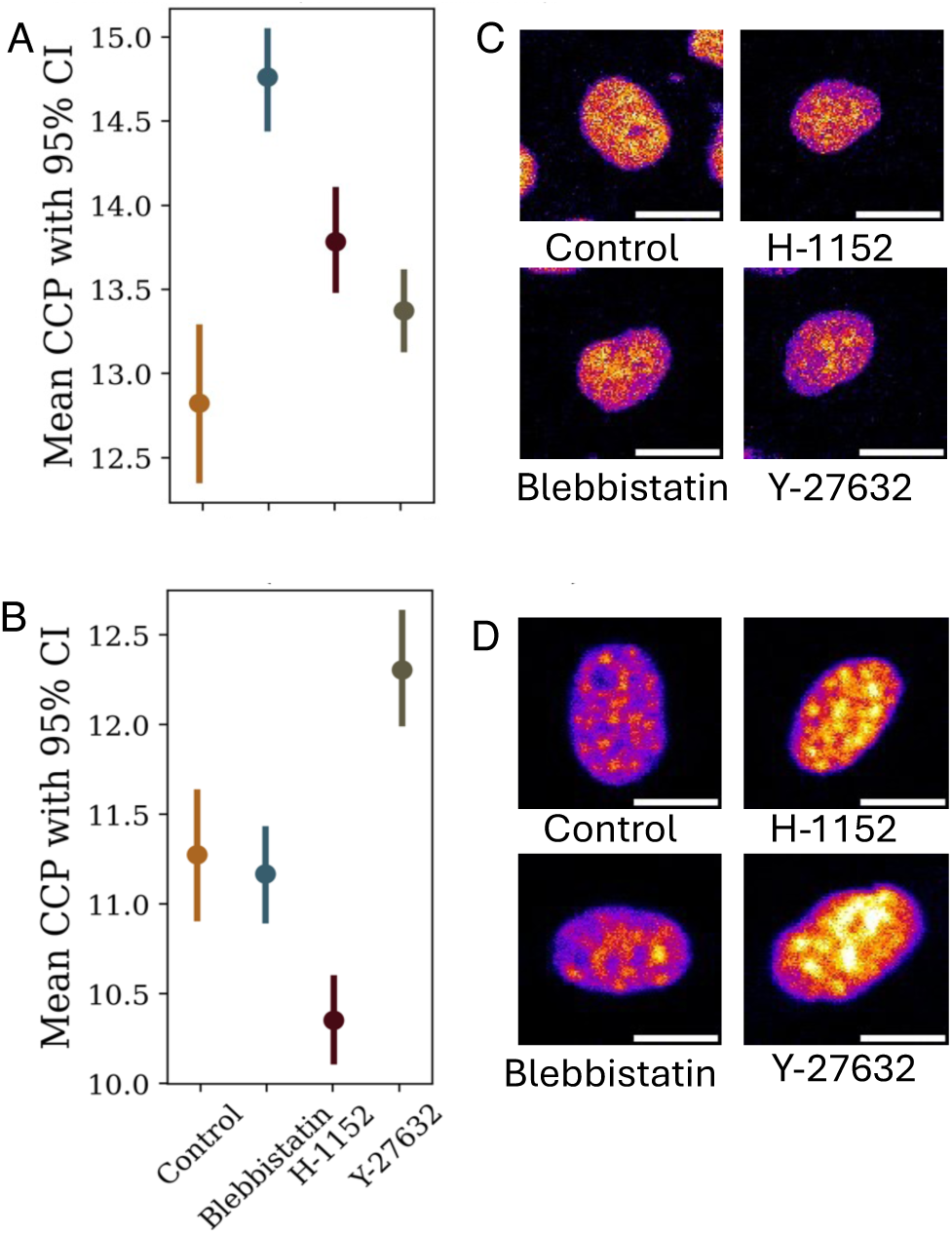
Chromatin organization by treatment. Mean Chromatin Condensation Parameter with 95% confidence intervals for nuclei in HeLa Cells (A) and NIH3T3 Cells (B) with error bars representing the 95% confidence intervals. Representative images of nuclei for HeLa cells (C) and NIH3T3 cells (D) for each treatment group.

## Discussion

The determination and maintenance of nuclear shape has been an active area of research in recent years. In this paper we take a novel approach: we study cell-nucleus coupling by data-driven modeling, specifically by identifying which features of nuclear shape are predictable from cell shape. We then perturb cells by disrupting actomyosin contractility and reconstruct quantitative models and then examine how they changed. Using a combination of machine learning techniques and standard statistical analysis, we show that there is a strong coupling between cell shape and nuclear shape. Specifically, we find that the size of the nucleus is very well predicted by cell shape parameters, and so are several other attributes of nuclear shape such as flatness and tilt. Previous papers have suggested that the nucleus could be actively shaped by contractile forces. To study whether nuclear shape and size were maintained by prestress created by actomyosin contractility, we treated HeLa cells with actomyosin contractility disrupting drugs. We show that relaxation of prestress due to actomyosin contractility does indeed affect the size of the nucleus but leaves the shape, i.e., features like elongation and solidity, largely unchanged. This is in accordance with other work that found, for example, that nuclear elongation does not change after nuclei are isolated from the cell ^41^. Cytoskeletal prestress thus is needed to maintain nuclear size but does not strongly affect other shape features at least in adherent HeLa cells. Nevertheless, many nuclear shape features are coupled to cell shape features as we have shown by the analysis of shape modes in hiPSCs. These could be due to geometric constraints arising from caging by vimentin or microtubules or the ER, for example. These factors could be fruitfully studied in future using our data-driven approach.

We also show that relaxation of prestress leads to changes in chromatin organization in a cell line specific way. In HeLa cells in particular this leads to an increase in chromatin compaction, while in NIH3T3 fibroblasts we see both increase and decrease. Non-parametric tests confirm that these changes are statistically reliable, and though they are small, it is important to note that these are global changes, and thus are likely to have functional consequences.

We find that machine learning models can predict nuclear shape with fairly high accuracy. It is certainly possible to increase the accuracy we report here by using more powerful models and by using more data. For the Cell2Nuc U-Net mode, our dataset for the hiPSCs was under 4,000 cells and for the HeLa cells was under 2,200 for each treatment. We found that the U-Net model needed retraining for the HeLa cells but converged rapidly when initialized with our pre-trained models on the hiPSC data. One of the challenges of the Cell2Nuc model was that we were attempting to predict a very large number of voxels without any constraints. It may be possible that a loss function combining the binary cross-entropy with a measure of accuracy such as mean squared error of the dimension calculations from the image could result in higher quality predictions. The ANNs using the shape modes performed very well and helped us realize that cell shape was informative of some of the more subtle features of nuclear shape. Since the shape modes are interpretable, we were able to understand some of our results at a deeper level. Currently, there is significant interest in applications of machine learning to biological questions. Our paper illustrates one approach to this problem, based on the capacity of machine learning models to function as universal approximators ^45^.

Our results have interesting implications for mechanical models of the nucleus. Previous literature has assumed that the nucleus can be modeled as either an elastic object or a liquid drop ^29,30^. Our finding that the shape of the nucleus is unchanged after abrogation of actomyosin contractility is in accordance with the nuclear drop model, but we find that its size changes at short ( ∼30 minutes) timescales in both HeLa and NIH3T3 cells. It’s not clear how this can be explained by either model, and raises interesting questions for future work.

One of the general conclusions of this study is that due to the mechanical and geometric constraints that the cell imposes on the nucleus, they are strongly coupled. Thus, it is possible to predict many features of nuclear shape from cell shape. This result advances one more reason why so many studies have found cell shape to be highly informative for cell state ^25,26,46–51^ -- it is also because cell shape by itself encodes major features of nuclear shape.

### Limitations of the Study

We used only a limited subset of just over 3800 images to train and test the Cell2Nuc UNet model. Computational limitations prevented using a larger set of images. We believe that this model could have performed even better with a larger training dataset. We were also unable to explore different machine learning architectures for this purpose, but plan to explore this in subsequent work. We abrogated actomyosin contractility in the HeLa cell and NIH3T3 cell study, but we did not perturb either the microtubule network or the vimentin network. Both have been left for later work.

## STAR Methods

### Allen Institute for Cell Science Database

The cell and nuclear images examined in this work come from the public database described in Viana et. al. ^33^. Their database contains 215,081 3-dimensional images of hiPSCs from the WTC-11 cell line which were genetically modified to produce fluorescent markers for various subcellular structures, such as actin filaments and microtubules. The cells were grown in colonies and imaged with spinning disk confocal microscopes with the DNA and cell membrane stained to yield 18,100 FOVs with the nucleus, cell membrane, structure of interest, and brightfield channels.

From their images, the nuclear lamina and cell membrane were segmented with machine learning models and manually validated as well as examined for outliers. A total of 578 spherical harmonic expansion (SHE) coefficients were calculated for the 3D shapes of these segmentations, truncated at order l = 16. These coefficients’ 3D representation was found to have a mean distance of 0.28 microns and 0.14 microns for the cell and nuclear membranes respectively. A principal component analysis (PCA) found the top 8 components to describe the variation in cell and nuclear shape from the SHE coefficients to create what the paper describes as ‘shape modes.’

All of this data, as well as the cell’s stage in the cell cycle, whether it was on the edge of the colony, whether it was marked as an outlier from artefacts in the segmentation process, and the calculated dimensions of height, surface area, and volume for the cell and nucleus, were reported in the metadata of the database. This work utilizes the 197,682 non-edge, non-outlier interphase cell samples from the reported metadata as well as the 3,824 non-outlier interphase cell samples with fluorescent markers for actin filaments from the collected images and segmentations.

### Allen Data Processing Steps

Before the data could be used to train a neural network model for prediction, each of the variables had to be scaled so that each contributed equal weight to the model’s performance. The height, surface area, volume, and each of the shape modes for the cell and nucleus were converted into z-scores for that variable, and the means and standard deviations for their distributions were recorded.

### Shape Mode Visualization

To create the shape mode visualization plots, the z-score scaling had to be reversed, as well as the PCA by multiplying the shape modes with their covariance matrix, returning values for all 289 SHE coefficients of the cell and nuclear membrane shape.

Applying these SHE coefficients and reconstructing the 3D image, a base spherical mesh of 10,000 points was created in spherical coordinates from a meshwork of 100 evenly spaced azimuth points and 100 evenly spaced elevation points. The associated Legendre polynomial values were calculated over this mesh, and the real form of the solid harmonics functions were used to add each harmonic layer weighted by the SHE coefficient. The coordinates were converted to Cartesian, and a final scaling factor of 0.108333 was applied to the voxels to represent distance in microns.

### Shape Reconstruction

To plot the reconstructed images described by the shape modes predicted by the models, the z-score scaling had to be reversed from these predictions with the previously recorded means and standard deviations of the shape mode data. As well, the PCA was reversed by multiplying the 12 shape modes with the covariance matrix, returning predictions for all 289 SHE coefficients of the cell and nuclear membrane shape. The same method of applying these SHE coefficients to the real solid harmonics functions on a 3D spherical mesh was used.

Images of the cell membrane were also imported to determine the rotation required to compare the predicted nuclear shape and true nuclear image. This was necessary since the SHE coefficients were all originally calculated with the longest cell dimension aligned to the x axis for standard procedure.

### Neural Network Architecture and Methods

The hidden layers of the ANN models included 3 layers with the same number of nodes as the largest of the input and output layers, with the largest model including 23 node hidden layers and the smallest including just 8. Each model was trained with Matlab’s Deep Learning Toolbox, and particularly the Bayesian regularization with backpropagation training function with the default parameters. Training included 10-fold cross validation with 85% of the total dataset for training and 15% for validation in each fold. The models were evaluated by the mean squared error of their predictions over 100 total epochs, after which the root mean squared error and correlations for each predicted variable were saved for the models and reported as averages across the 10 folds. All models were trained on a 10 core i9 processor with boosted 4.5 GHz speed and minimal RAM use and took roughly a few days to run the full cross-fold validation.

### Multiple Linear Regression Models

Each of the MLR models was created with Matlab’s stepwise linear regression function with a p-value cutoff of 10-20 (adjusted for sample size) and multiplicative interactions allowed between the input variables to provide some non-linearity in the model. The same as for the ANN models, the RMSE and correlations were collected for each of the predicted output variables. All models were similarly trained on a 10 core i9 processor with boosted 4.5 GHz speed and minimal RAM use and took roughly a few hours to run the MLR models each.

### Decomposition of Total Variance in MLR Models

Individual contributions to the total explained variance of the MLR models by any single cell membrane predictor were estimated via the following assumptions, prerequisites, and equations. It was assumed that each of the predictor variables were not collinear with any other, this being justifiable for the cell shape modes as they are the result of PCA and therefore define orthogonal principal components. Each of the input predictor variables were normalized and scaled such that their mean was 0 and standard deviation was 1. And the following equations were used to define these individual contributions, or percent explained variances, for each predictor variable, where R_*i*_^2^ represents the percent explained variance of the ith predictor, *β̂*_*i*_ represents the slope coefficient of the ith predictor in our model, COV() represents a function calculating the covariance of two variables, *x*_*i*_ represents the ith predictor, and *y*’ represents the nuclear shape parameter predictions made by the model:

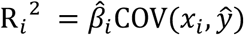

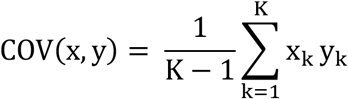

### 3D U-Net Generative Prediction Models

A diagram of the U-Net architecture of this model can be seen in Supplementary Figure S2 and was based on the work of Cirillo et. al. [5]. The model consists of a series of 6 3D convolution steps with a kernel size of 4×4×4 and stride 2, followed by instance normalization and a leaky ReLU activation function, as well as a series of 6 3D transpose convolution steps with the same kernel size and varying stride, cropping, and padding to allow for skip connections between the downward and upward convolutional steps. The final convolution applies a sigmoid activation function to limit the predicted nuclear image to output voxel values between 0 and 1 which can be interpreted as a log probability that a particular voxel is part of the nuclear mask image.

The model was created in TensorFlow and trained in batches of 10 images on an Nvidia A100-SXM4-80GB GPU over 20 epochs, each taking about 3 days to complete. The BCE reconstruction loss function was chosen alongside the Adam optimizer with a learning rate of 0.0002 and default parameters aside from choosing an exponential decay rate of the first moments beta 1 to be 0.5. As a measure to prevent overfitting, the input membrane images in each batch were randomly chosen to be augmented by either a reflection or rotation, or a combination of the two.

### Interpreting U-Net Predictions

The predictions yielded from the cross-validated U-Net models were interpreted as confidence values in a particular voxel belonging to the nucleus image since the final sigmoid activation layer produces values in the continuous range from 0 to 1. Therefore, it was necessary to employ a threshold value below which the confidence is considered too low to include as part of the nucleus prediction so that the prediction could be interpreted as a binary image of only background and nucleus voxels. It should be noted that the resulting data measured from the prediction with the threshold applied was quite sensitive to this threshold value, and efforts were taken to choose the best threshold for the set of 10 cross-validated models by assessing these results over a large range of threshold values and selecting the value which maximized the models’ accuracy, finding 0.42 to be this value for the Allen image dataset and 0.55 for the HeLa image dataset. In hindsight, this threshold value could have been included in the model itself as a hyperparameter and applied to the prediction before the calculation of BCE loss to avoid this search for a “best value”.

These binary prediction images were then used in calculating Dice scores as well as measuring nuclear dimensions like height, surface area, volume, and tilt. Particularly in the case of surface area, but applied to all measures, a median filter (of size 3×3×3 voxels) was deemed necessary to smooth out the binary image since the model had not been trained with the threshold as a hyperparameter, leading to binary images with rough edges at the threshold.

### Measuring Dimensional Data from U-Net Binary Predictions

The height was measured from the binary prediction images by finding the distance between the maximum and minimum z-stacks of the predicted nucleus and multiplying by a scaling factor of 0.29 µm for the Allen database cells and 0.41 µm for the HeLa set. The surface area was calculated with a 3×3×3 voxel convolution with stride 1 and same padding such that the exposed faces of the voxels at the surface of the nucleus were assigned a value corresponding to the actual area of that face in µm² and later summed. The volume was simply calculated by summing the number of voxels the nucleus contained and multiplying by the appropriate scaling factor of 0.0034 µm³ for the Allen set and 0.0176 µm³ for the HeLa cells. A measure of nuclear tilt was achieved by finding the center of the maximum and minimum z-stacks and calculating their distance divided by the earlier calculated height. The angle Θ created by this axis of tilt and the z axis was then calculated by applying arccosine to the reciprocal of this distance value.

### Shape Parameters Calculated with Dimensional Data from U-Net Predictions

With the height, surface area, and volume represented as H, A, and V respectively, the following equations describe the measure (S) of how spherical a particular nucleus was, given the radii of equivalent spheres for each of the dimensions:

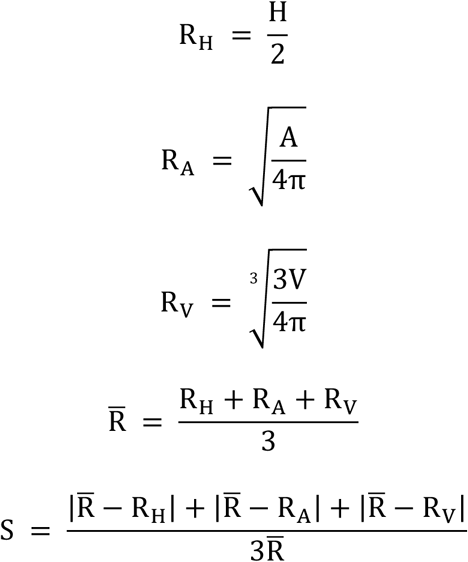

In addition, a flattening parameter (F) describing the height of a nucleus as compared to the radius of a sphere equivalent to the nucleus volume was calculated with the following equation:

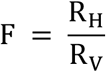

### Cell Culturing and Preparation

HeLa cells (Kyoto line transformed with Myosin-II-GFP as validated by Dr. O’Neil Wiggan) were generously gifted by Dr. Diego Krapf. NIH3T3 fibroblast cells were generously gifted by Dr. Soham Ghosh. The culture conditions for both cell lines were 5% CO2, 37°C, 90% relative humidity. Cells were maintained in HyClone Dulbecco’s Modified Eagle Medium (DMEM), F12 with L-glutamine, HEPES (Global Life Sciences) supplemented with 10% EqualFetal bovine serum (Atlas Biologicals) and 1% pen/strep (Quality Biological) and passaged at 80-90% confluency after trypsinization with TrypLE Express (Gibco).

Pharmacological treatment: Live cells were treated with 0.3µM H-1152 dihydrochloride (Enzo Life Science), 10µM Y-27632 (Tocris) or 0.1µM para-amino-blebbistatin (Cayman) in DMEM for 30 minutes at 37°C prior to fixing. Control group was maintained in DMEM prior to fixing.

Cell staining: For all experiments cells were seeded on Mattek 35mm round glass bottom imaging dishes at concentrations of 15,000-25,000 cells/ml. Cells were fixed with 3.7% Paraformaldehyde (PFA, Thermo Scientific) (in 1X PBS) for 10 minutes at room temperature. Cells were permeabilized in 0.1% Triton X-100 in 1X PBS. Normal goat serum (Jackson Labs) and bovine serum albumin (Sigma-Aldrich) was used for blocking for non-specific binding. Cells were stained for F-actin with Phalloidin 594 (Invitrogen, 400x) at a dilution of 2.5:1000 and for chromatin with 4’,6-diamidino-2-phenylindole (DAPI, 300µM stock solution) at a dilution of 1:1000.

### Imaging

Fluorescent images were acquired with Zeiss laser scanning confocal microscope with 40x/1.2NA water objective. Images are z-stacks of 20-60 slices with step size of 0.42 µm, captured using Zen Blue software. Lasers included the excitation wavelength of 405nm, 488nm and 561nm. All imaging were performed at 3% laser power.

### Image processing and Morphological Feature extraction

2D Image processing: Maximum projections of multichannel z-stack images were segmented using CellProfiler. Nuclei were identified by the DAPI channel followed by cell identification using the myosin-GFP channel. Morphometric data for each identified cell object and nucleus object was extracted. Additional nuclear and cell parameters were calculated using custom python scripts. Nuclear images were also segmented separately in CellProfiler prior to CCP analysis.

3D Image processing: multichannel z-stack images were segmented using Cellpose within the framework of CellProfiler. Prior to segmentation, images were trimmed in ImageJ FIJI. The trimming consisted of deleting layers where edge detection and variance were below a threshold. These trimmed images were then fed into Cellprofiler where a Cellpose plugin identified the cell outlines to create masks. The method consisted of identifying cells and creating a mask for each z-plane then stitching these layers together to reconstruct the 3D object. These masks were then filtered via volume and position, to remove unreasonable Cellpose predictions and remove cells touching the image edges. These cell masks and nuclear masks were then paired via the relate objects module in Cellprofiler. Morphometric data for each identified cell object and nucleus object was extracted.

### 3D HeLa Cell Data Processing Steps and Model Training

The HeLa cell and nuclear masks were manually validated and examined for outliers with a simple measure of ensuring that the bounding boxes of the nuclei fit entirely within that of the cell as well as analysis of histograms of the dimensional data extracted from these nuclear masks. With these methods, the sample size of cell and nuclear mask pairs for the different groups were reduced to 1072, 2148, 1214, and 1413 for the control, blebbistatin, Y-27632, and H-1152 treatments respectively.

The U-Net models were retrained with this dataset for each treatment individually, with the same 10 cross-fold validations, architecture, hyperparameters, and training process, but only 10 epochs were trained since the models were given the 7^th^ U-Net model weights from the Allen dataset as a starting point. This model was close to the average performance of the cross-folds and therefore deemed appropriate for transfer learning.

### 2D HeLa and NIH3T3 cell data processing and analysis

Custom python scripts were written to process and analyze 2D morphological features extracted from maximum projection images of HeLa and NIH3T3 cells. Additional morphological metrics were calculated including Aspect Ratio, Circularity and Eccentricity. Outliers relative to cell and nuclear area were removed using Interquartile Range (IQR) with a multiplier (k) of 1.5 for nuclear area and 2.0 for cell area for HeLa cells and 1.0 for nuclear area and 1.5 for cell area for NIH3T3 cells. The first quartile (Q1) represents the 25^th^ percentile of the data, while the third quartile (Q3) represents the 75^th^ percentile for the selected columns. The IQR is calculated as:

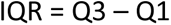

This captures the middle 50% of the data. Outliers are identified as points lying outside this range +-the multiplier (k) as follows:

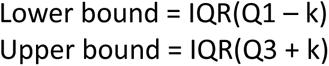

For the HeLa cell type, a total of 312 identified cells out of 6124 total were removed. Of the remaining cells the control group was represented by 975, 1240 for the H-1152 group, 1458 for the Y-27632 group and 2139 in the Blebbistatin group. For the NIH3T3 cell type, a total of 215 identified cells out of 1655 total were removed. Of the remaining cells the control group was represented by 241, 352 for the H-1152 group, 539 for the Y-27632 group and 308 in the Blebbistatin group. Summary statistics and Histograms of size and shape features for both cell and nucleus were examined prior to correlation and regression analyses.

A publicly available MATLAB script was used to quantify chromatin condensation ^44^. Individual nuclear maximum projection images were first standardized via intensity profile redistribution. Sobel edge detection was applied to construct a new edged image based on the gradient magnitude. Next the image is thresholded to determine strong edges followed by thinning to reduce elements to a single pixel width. Finally, the edge detected corresponding with the true edge of the nucleus is removed as it does not represent chromatin condensation. The resulting number of edges measured is divided by the area of the nucleus to result in the CCP. Nuclei counts for HeLa cells per treatment group are: 681 blebbistatin; 371 control; 773 H-1152; 1128 Y-27632. Nuclei counts forNIH3T3 cells per treatment group are: 468 blebbistatin; 375 control; 763 H-1152; 405 Y-27632.

## Resource Availability

### Lead Contact

Ashok Prasad

### Materials Availability

This study did not generate any unique reagents or other materials.

## Data and Code Availability

Images from the Allen Institute hiPSC cell image database can be accessed from this page: https://www.allencell.org/data-downloading.html

Images of cells taken for this study and the trained models generated have been deposited to a repository in DataDryad and can be accessed at the following permanent identifier after publication of the manuscript: https://doi.org/10.5061/dryad.zgmsbccp1

All original code has been deposited at Github and is publicly available at https://github.com/prasadlabcsu/NucleusPrediction as of the date of publication.

A copy of all of the code has also been placed in the DataDryad repository for permanent archiving.

## Acknowledgements

This work was supported by an NSF BRITE award CMMI-2227605 to AP and a NIH T32 Fellowship (Award No: T32GM132057) to RD.

## Author Contributions

Conceptualization: A.P., S.L and R.D; Methodology: A.P., S.L. and R.D.; Investigation: A.P., S.L., R.D., R.S., S.G. and S.S.; Writing—original draft: A.P., S.L. and R.D.; Writing—review & editing: all authors; Funding acquisition: A.P.; Resources: S.G., O.W.; Supervision, A.P.

## Declaration of Interests

All authors declare that they have no conflicts of interest.

## Notes

### Competing Interest Statement

The authors have declared no competing interest.

