## Supplemental Information for "Cell Morphology accurately predicts the nuclear shape of adherent cells"

**Contents:**

**Figures S1-S6**

**Tables S1-S4**

|  | **Percent Variance Captured by PCA of Cell and Nuclear Shape Modes** | | | | | | | |
| --- | --- | --- | --- | --- | --- | --- | --- | --- |
| **Shape Mode** | |  | **Cell** | **Cell Cumulative Total** |  |  | **Nucleus** | **Nucleus Cumulative Total** |
| 1 | |  | 19.88 | 19.88 |  |  | 35.94 | 35.94 |
| 2 | |  | 12.58 | 32.46 |  |  | 19.67 | 55.61 |
| 3 | |  | 11.91 | 44.37 |  |  | 9.18 | 64.79 |
| 4 | |  | 7.49 | 51.86 |  |  | 7.01 | 71.80 |
| 5 | |  | 6.76 | 58.62 |  |  | 6.78 | 78.58 |
| 6 | |  | 5.31 | 63.93 |  |  | 6.26 | 84.84 |
| 7 | |  | 4.97 | 68.90 |  |  | 1.37 | 86.21 |
| 8 | |  | 2.30 | 71.20 |  |  | 1.26 | 87.47 |
| 9 | |  | 1.98 | 73.18 |  |  | 1.12 | 88.59 |
| 10 | |  | 1.77 | 74.95 |  |  | 1.04 | 89.63 |
| 11 | |  | 1.74 | 76.69 |  |  | 0.99 | 90.62 |
| 12 | |  | 1.45 | 78.14 |  |  | 0.92 | 91.54 |
| 13 | |  | 1.24 | 79.38 |  |  |  |  |
| 14 | |  | 1.22 | 80.6 |  |  |  |  |
| 15 | |  | 1.19 | 81.79 |  |  |  |  |
| 16 | |  | 1.10 | 82.89 |  |  |  |  |
| 17 | |  | 0.98 | 83.87 |  |  |  |  |
| 18 | |  | 0.84 | 84.71 |  |  |  |  |
| 19 | |  | 0.83 | 85.54 |  |  |  |  |
| 20 | |  | 0.76 | 86.30 |  |  |  |  |

**Table S1:** Table of the percent variance captured by each cell and nuclear shape mode form the principal component analysis of the spherical harmonic expansion coefficients.

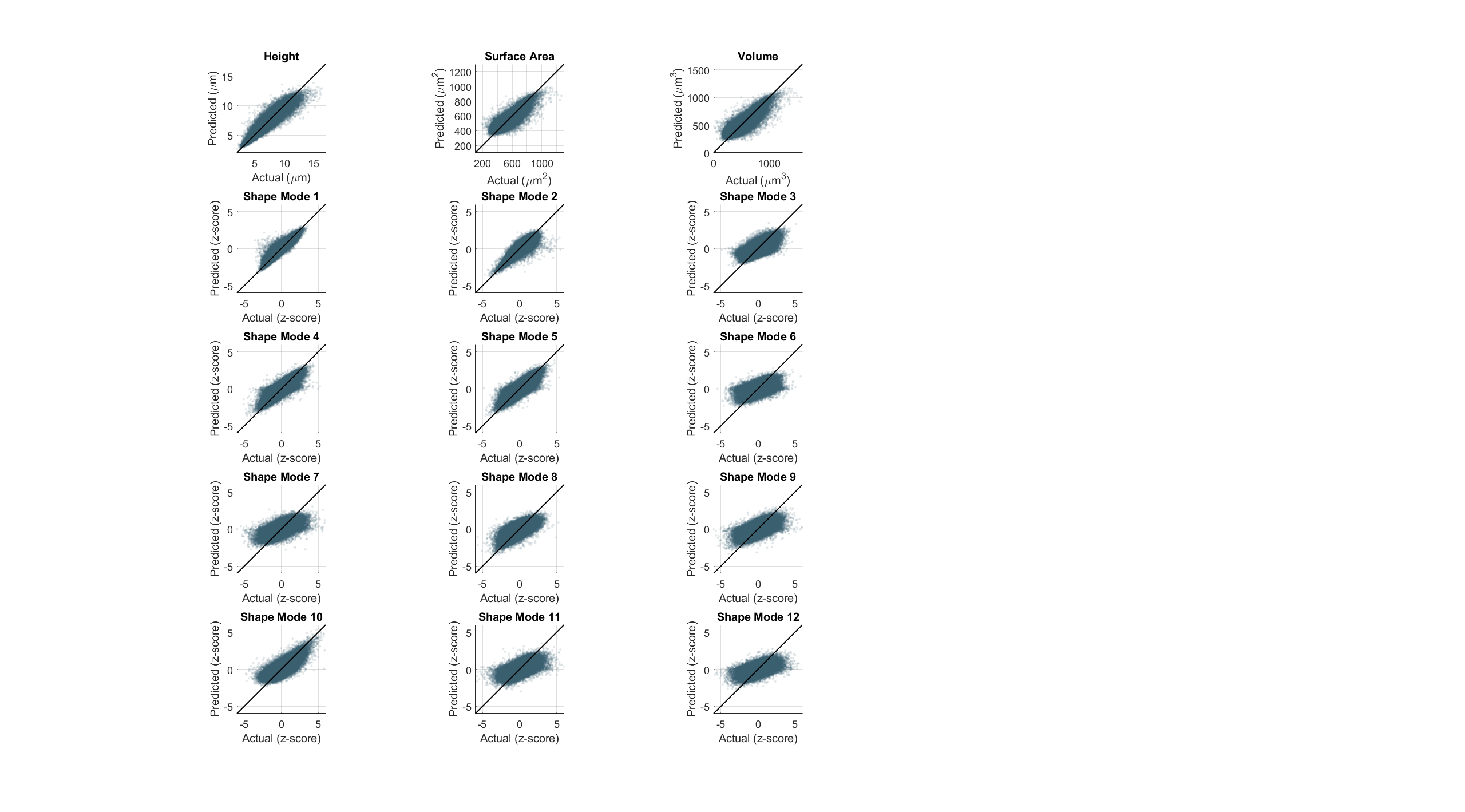

**Figure S1.** Scatter plots of predicted vs actual values for the nuclear shape parameters. Every point is a cell. Note that the values of the shape parameters have been normalized by subtracting the mean and dividing by the standard deviation (referred to here as z-score).

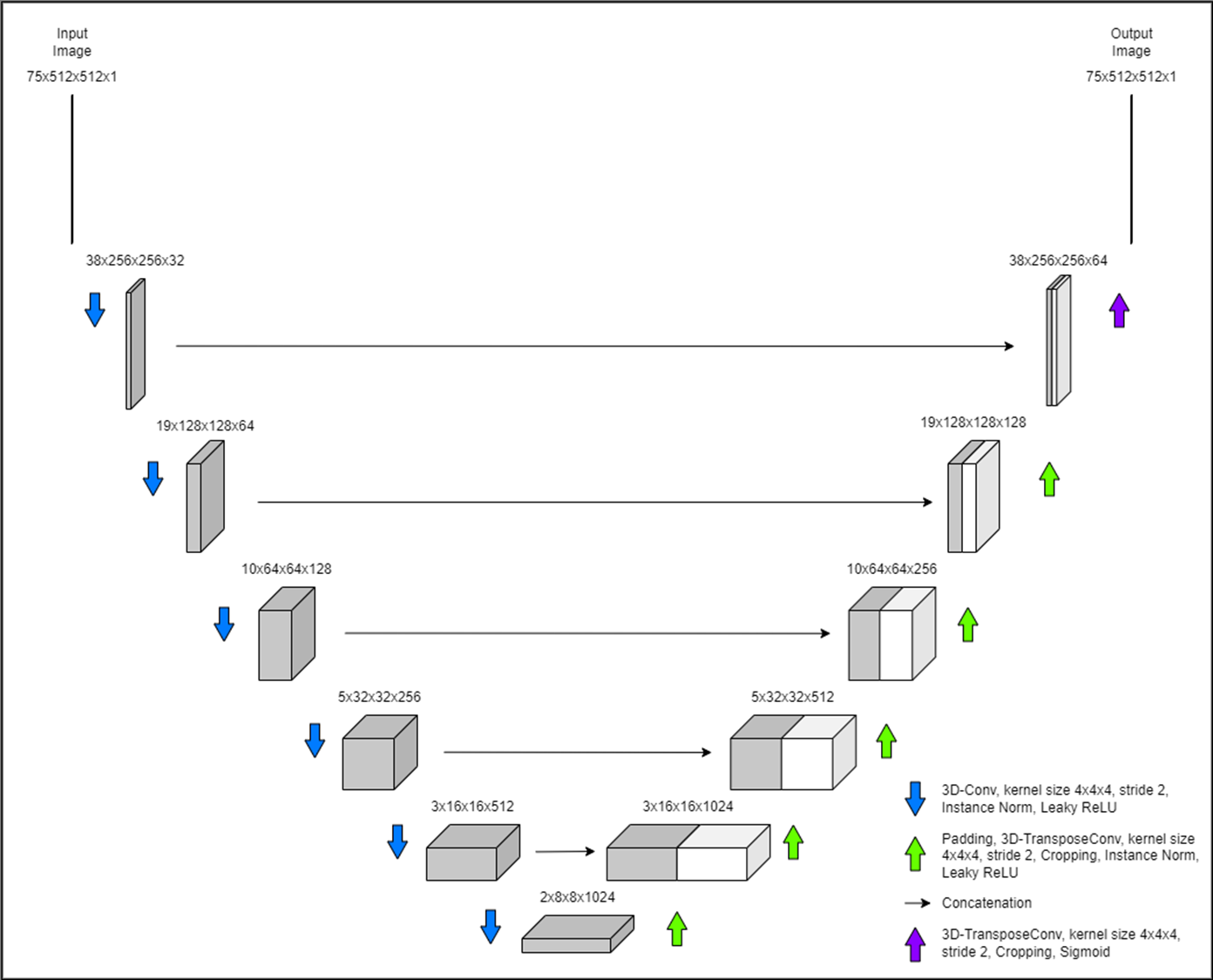

**Figure S2.** Diagram of the U-Net architecture utilized in the convolutional neural network models with the dimensions of the resulting images at each step and labels corresponding to the operations applied in each step.

| **Model:** | **Includes Cell Height, Area, and Volume** | | | **Excludes ell Height, Area, and Volume** | | | | **Includes Only Shape Modes 1-5** | | | |
| --- | --- | --- | --- | --- | --- | --- | --- | --- | --- | --- | --- |
| **Nuclear Output:** | **Height** | | | | | | | | | | |
| **Input Variable** | **% EV** | **Total % EV** | **Total R²** | **Input Variable** | **% EV** | **Total % EV** | **Total R²** | **Input Variable** | **% EV** | **Total % EV** | **Total R²** |
| Shape Mode 1 | 35.9 | 85.4 | 88.8 | Shape Mode 1 | 74.4 | 78.5 | 80.5 | Shape Mode 1 | 74.8 | 78.5 | 78.6 |
| Height | 25.9 |  |  | Shape Mode 5 | 2.8 |  |  | Shape Mode 5 | 2.9 |  |  |
| Volume | 11.2 |  |  | Interactions | 1.3 |  |  | Shape Mode 3 | 0.8 |  |  |
| Surface Area | 6.4 |  |  | - | - |  |  | - | - |  |  |
| Interactions | 6.0 |  |  | - | - |  |  | - | - |  |  |
| **Nuclear Output:** | **Surface Area** | | | | | | | | | | |
| **Input Variable** | **% EV** | **Total % EV** | **Total R²** | **Input Variable** | **% EV** | **Total % EV** | **Total R²** | **Input Variable** | **% EV** | **Total % EV** | **Total R²** |
| Surface Area | 43.6 | 79.3 | 81.0 | Shape Mode 3 | 48.2 | 63.5 | 64.4 | Shape Mode 3 | 46.6 | 60.3 | 60.7 |
| Volume | 22.4 |  |  | Shape Mode 5 | 14.1 |  |  | Shape Mode 5 | 13.4 |  |  |
| Shape Mode 3 | 5.5 |  |  | Shape Mode 15 | 1.2 |  |  | Shape Mode 1 | 0.3 |  |  |
| Interactions | 5.1 |  |  | - | - |  |  | - | - |  |  |
| Height | 2.7 |  |  | - | - |  |  | - | - |  |  |
| **Nuclear Output:** | **Volume** | | | | | | | | | | |
| **Input Variable** | **% EV** | **Total % EV** | **Total R²** | **Input Variable** | **% EV** | **Total % EV** | **Total R²** | **Input Variable** | **% EV** | **Total % EV** | **Total R²** |
| Volume | 50.9 | 83.1 | 86.4 | Shape Mode 3 | 53.8 | 71.8 | 72.9 | Shape Mode 3 | 52.5 | 70.5 | 71.0 |
| Surface Area | 14.2 |  |  | Shape Mode 5 | 13.5 |  |  | Shape Mode 5 | 13.1 |  |  |
| Interactions | 12.6 |  |  | Shape Mode 1 | 4.5 |  |  | Shape Mode 1 | 4.9 |  |  |
| Height | 3.1 |  |  | - | - |  |  | - | - |  |  |
| Shape Mode 1 | 2.3 |  |  | - | - |  |  | - | - |  |  |
| **Nuclear Output:** | **Shape Mode 1** | | | | | | | | | | |
| **Input Variable** | **% EV** | **Total % EV** | **Total R²** | **Input Variable** | **% EV** | **Total % EV** | **Total R²** | **Input Variable** | **% EV** | **Total % EV** | **Total R²** |
| Shape Mode 1 | 37.4 | 82.2 | 87.4 | Shape Mode 1 | 69 | 81.0 | 82.0 | Shape Mode 1 | 69 | 81.1 | 81.2 |
| Volume | 23.0 |  |  | Shape Mode 3 | 7.3 |  |  | Shape Mode 3 | 7.3 |  |  |
| Interactions | 11.6 |  |  | Shape Mode 5 | 4.7 |  |  | Shape Mode 5 | 4.8 |  |  |
| Height | 5.7 |  |  | - | - |  |  | - | - |  |  |
| Shape Mode 3 | 4.5 |  |  | - | - |  |  | - | - |  |  |
| **Nuclear Output:** | **Shape Mode 2** | | | | | | | | | | |
| **Input Variable** | **% EV** | **Total % EV** | **Total R²** | **Input Variable** | **% EV** | **Total % EV** | **Total R²** | **Input Variable** | **% EV** | **Total % EV** | **Total R²** |
| Volume | 40.1 | 74.8 | 82.6 | Shape Mode 3 | 59.9 | 76.7 | 77.6 | Shape Mode 3 | 59.2 | 76.5 | 77.1 |
| Shape Mode 3 | 11.0 |  |  | Shape Mode 1 | 14.3 |  |  | Shape Mode 1 | 14.8 |  |  |
| Shape Mode 1 | 10.2 |  |  | Shape Mode 5 | 2.5 |  |  | Shape Mode 5 | 2.5 |  |  |
| Height | 6.9 |  |  | - | - |  |  | - | - |  |  |
| Interactions | 6.6 |  |  | - | - |  |  | - | - |  |  |
| **Nuclear Output:** | **Shape Mode 3** | | | | | | | | | | |
| **Input Variable** | **% EV** | **Total % EV** | **Total R²** | **Input Variable** | **% EV** | **Total % EV** | **Total R²** | **Input Variable** | **% EV** | **Total % EV** | **Total R²** |
| Shape Mode 5 | 21.9 | 39.4 | 40.8 | Shape Mode 5 | 34.7 | 36.6 | 38.5 | Shape Mode 5 | 34.5 | 36.2 | 36.4 |
| Interactions | 10.0 |  |  | Shape Mode 3 | 1.2 |  |  | Shape Mode 3 | 1.2 |  |  |
| Volume | 3.7 |  |  | Interactions | 0.7 |  |  | Shape Mode 1 | 0.5 |  |  |
| Surface Area | 2.8 |  |  | - | - |  |  | - | - |  |  |
| Shape Mode 3 | 1.0 |  |  | - | - |  |  | - | - |  |  |
| **Nuclear Output:** | **Shape Mode 4** | | | | | | | | | | |
| **Input Variable** | **% EV** | **Total % EV** | **Total R²** | **Input Variable** | **% EV** | **Total % EV** | **Total R²** | **Input Variable** | **% EV** | **Total % EV** | **Total R²** |
| Shape Mode 2 | 56.8 | 61.9 | 62.3 | Shape Mode 2 | 58.7 | 63.1 | 64.3 | Shape Mode 2 | 55.8 | 60.4 | 60.6 |
| Shape Mode 4 | 3.6 |  |  | Shape Mode 4 | 3.5 |  |  | Shape Mode 4 | 3.6 |  |  |
| Shape Mode 7 | 1.1 |  |  | Shape Mode 7 | 0.9 |  |  | Interactions | 1 |  |  |
| Shape Mode 18 | 0.2 |  |  | - | - |  |  | - | - |  |  |
| Interactions | 0.2 |  |  | - | - |  |  | - | - |  |  |
| **Nuclear Output:** | **Shape Mode 5** | | | | | | | | | | |
| **Input Variable** | **% EV** | **Total % EV** | **Total R²** | **Input Variable** | **% EV** | **Total % EV** | **Total R²** | **Input Variable** | **% EV** | **Total % EV** | **Total R²** |
| Shape Mode 4 | 55.3 | 60.4 | 60.9 | Shape Mode 4 | 55.3 | 59.6 | 60.9 | Shape Mode 4 | 55.3 | 58.9 | 59.1 |
| Shape Mode 2 | 3.5 |  |  | Shape Mode 2 | 3.5 |  |  | Shape Mode 2 | 3.5 |  |  |
| Shape Mode 6 | 0.8 |  |  | Shape Mode 6 | 0.8 |  |  | Shape Mode 5 | 0.1 |  |  |
| Shape Mode 20 | 0.6 |  |  | - | - |  |  | - | - |  |  |
| Shape Mode 9 | 0.2 |  |  | - | - |  |  | - | - |  |  |

**Table S2.** Percent explained variance of the top 5 contributors from the input variables of the multiple linear regression models that predict nuclear shape modes from cell shape modes and other features.

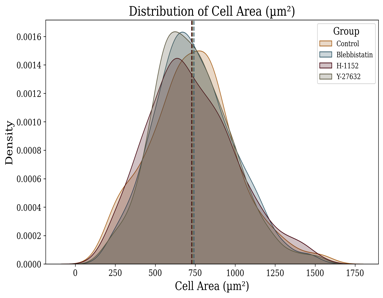

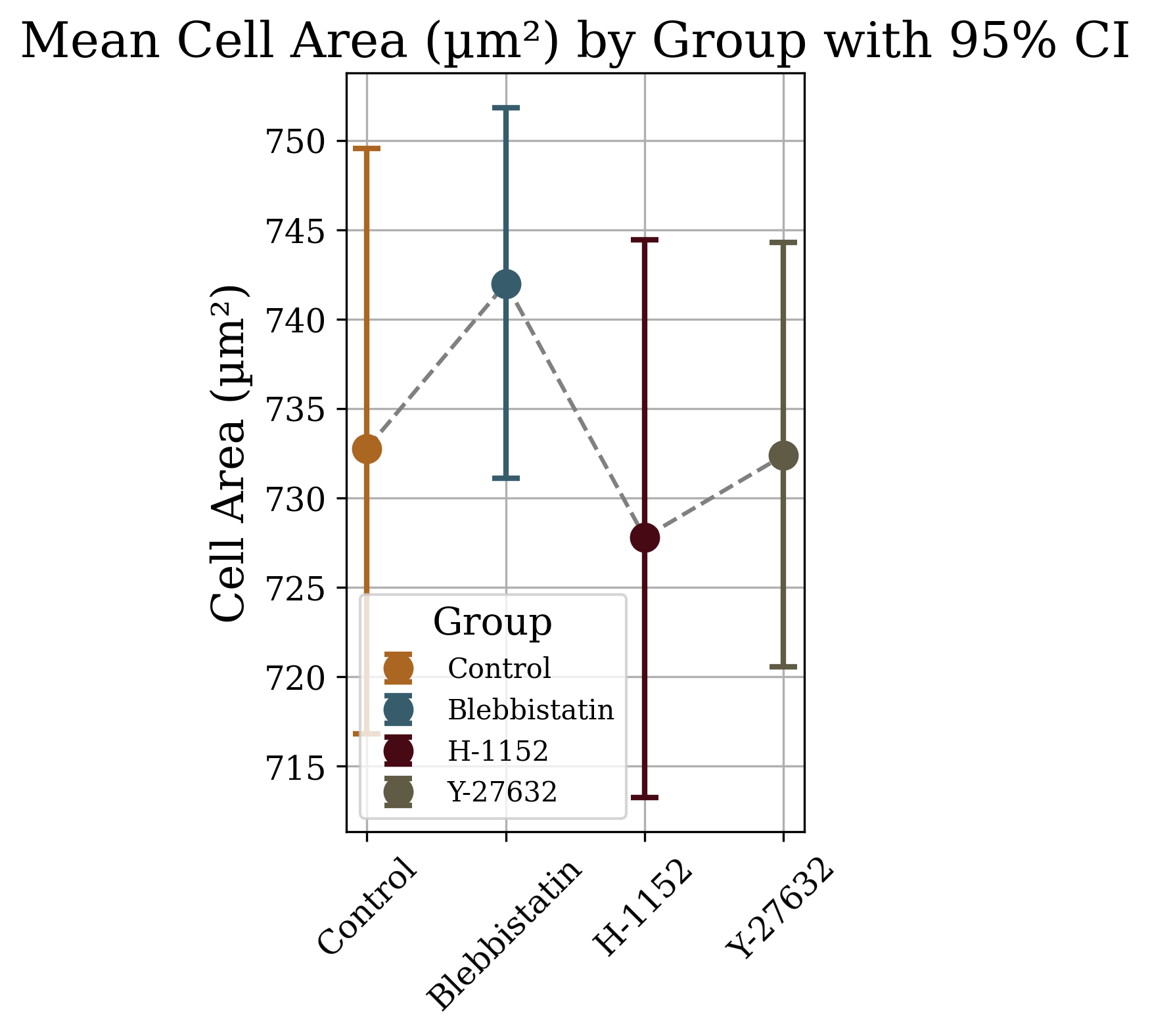

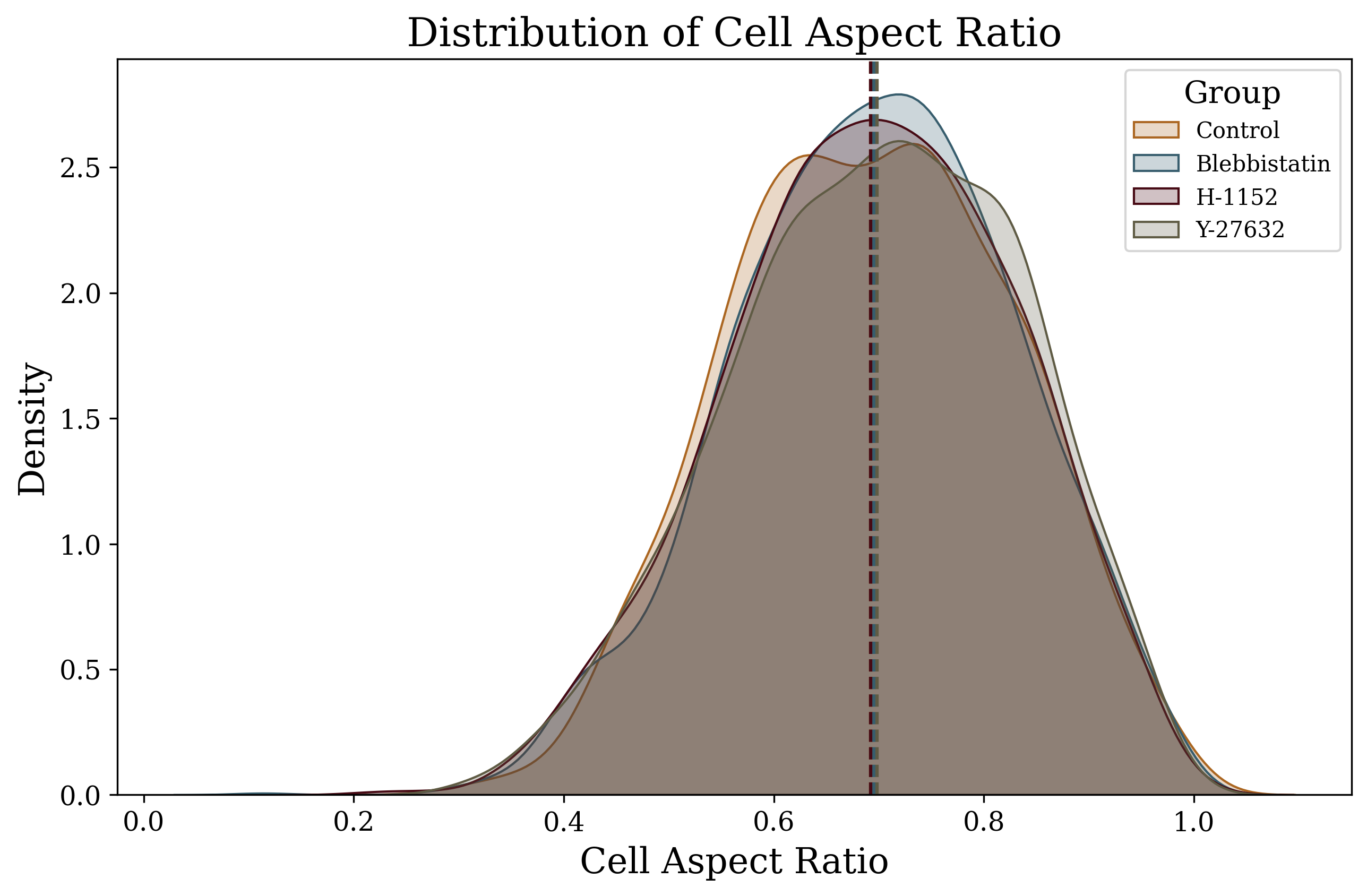

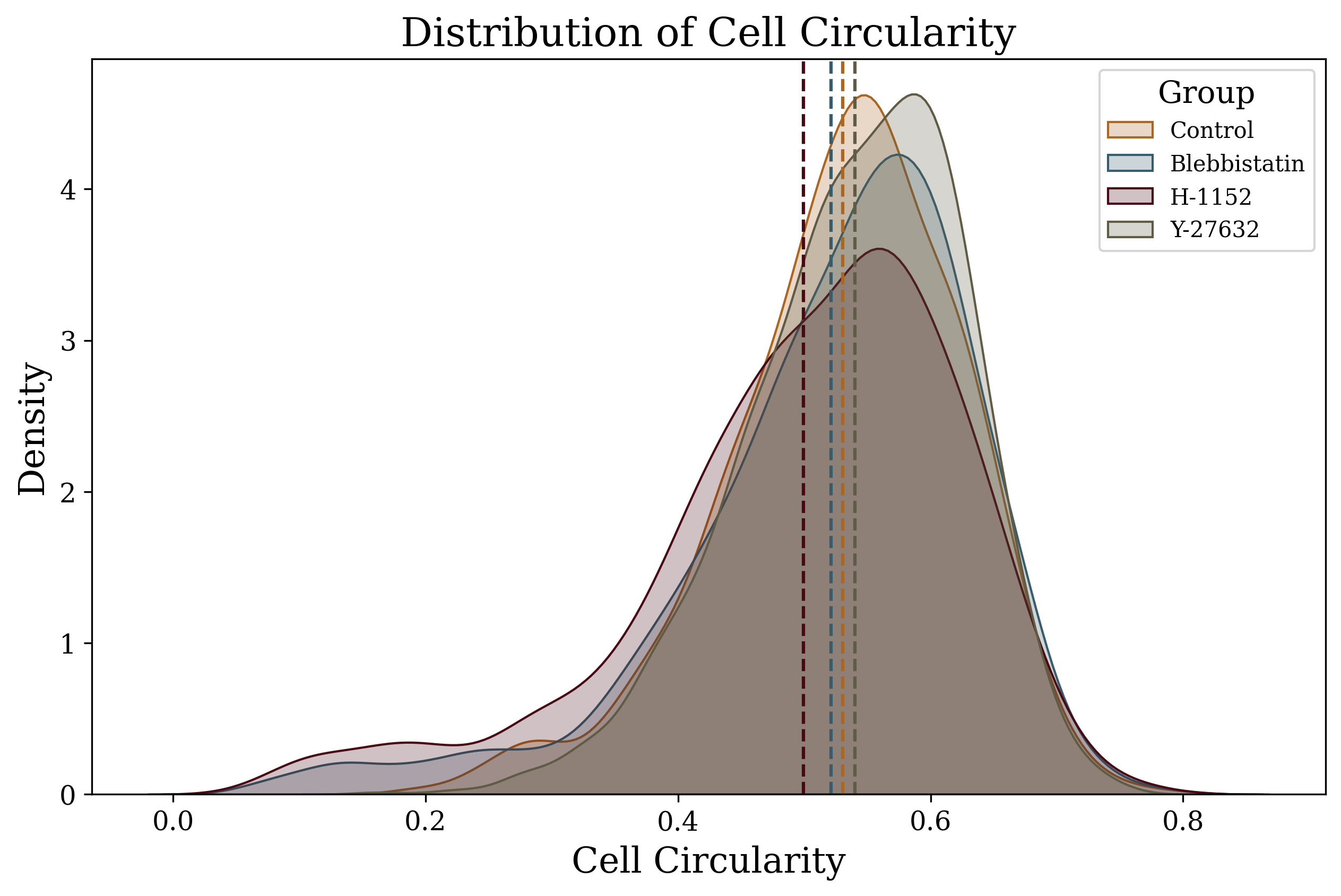

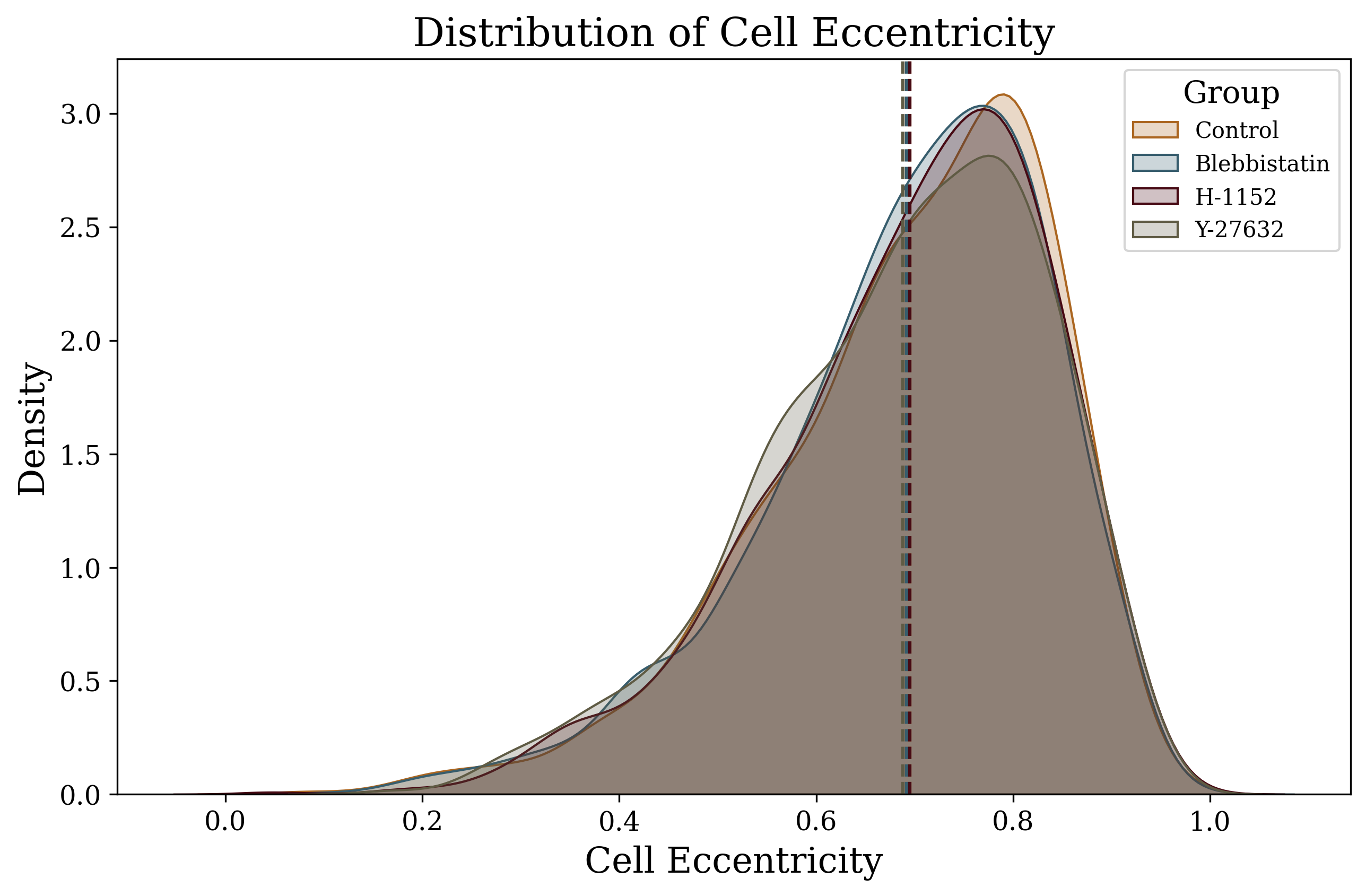

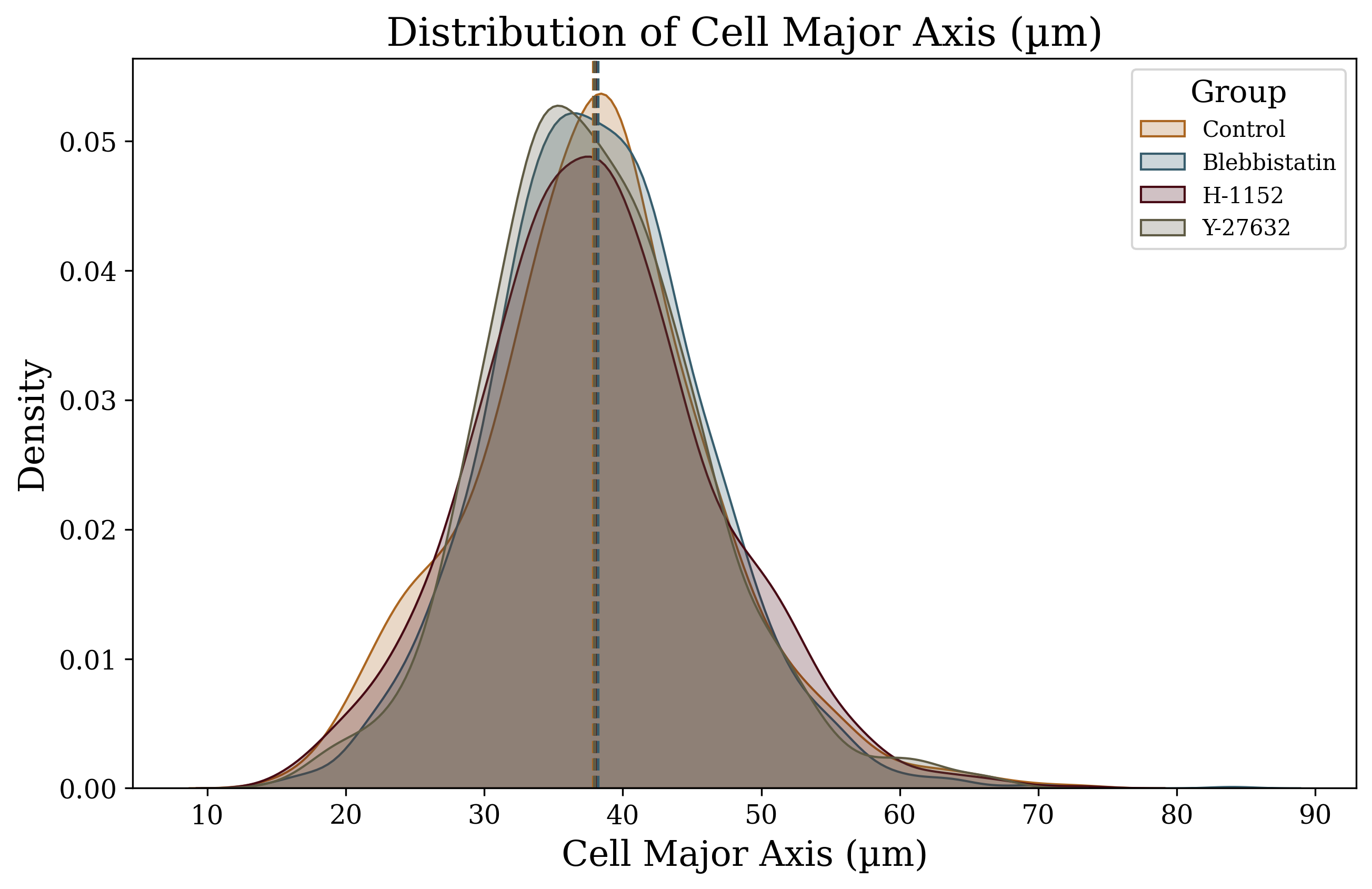

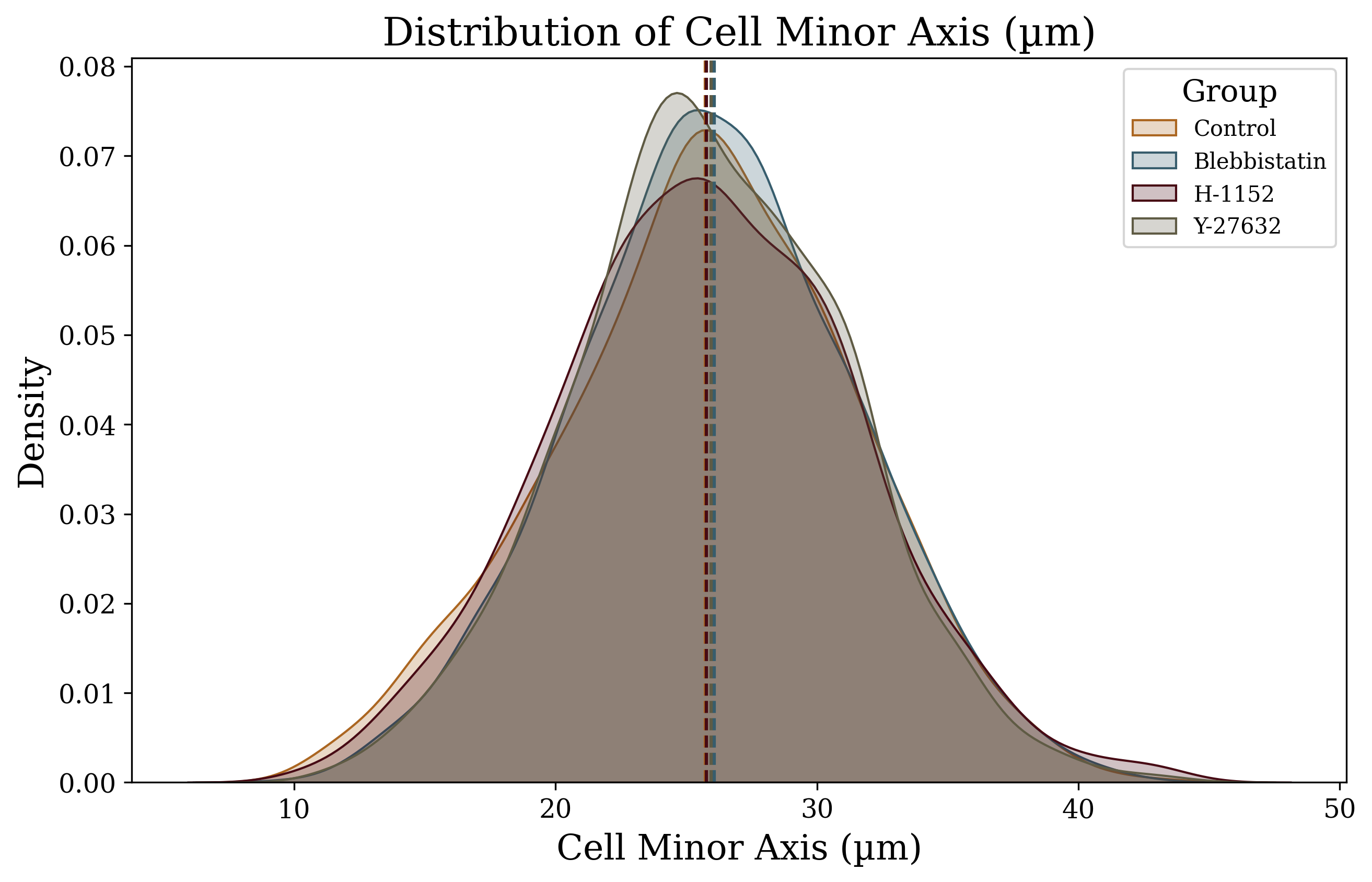

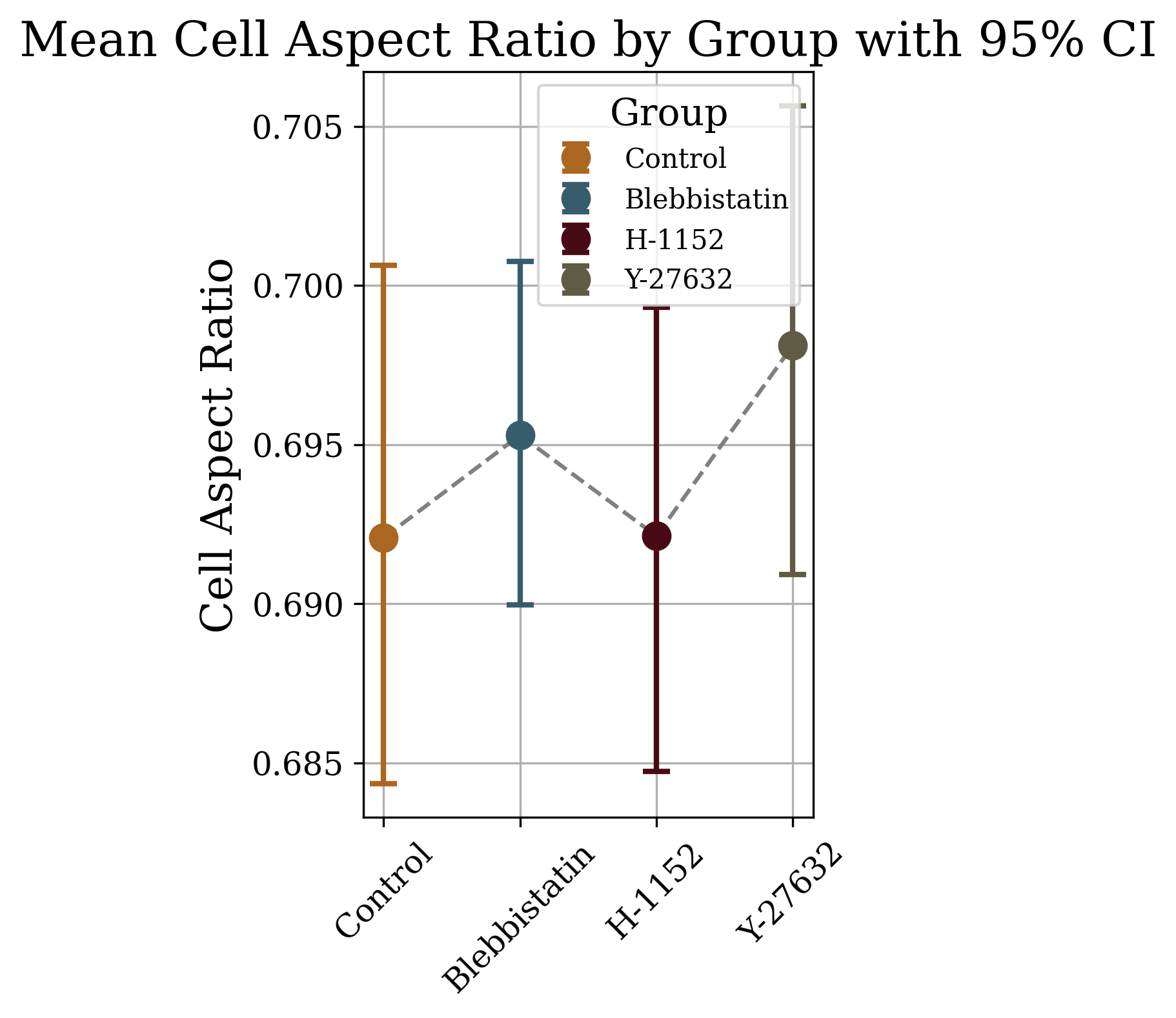

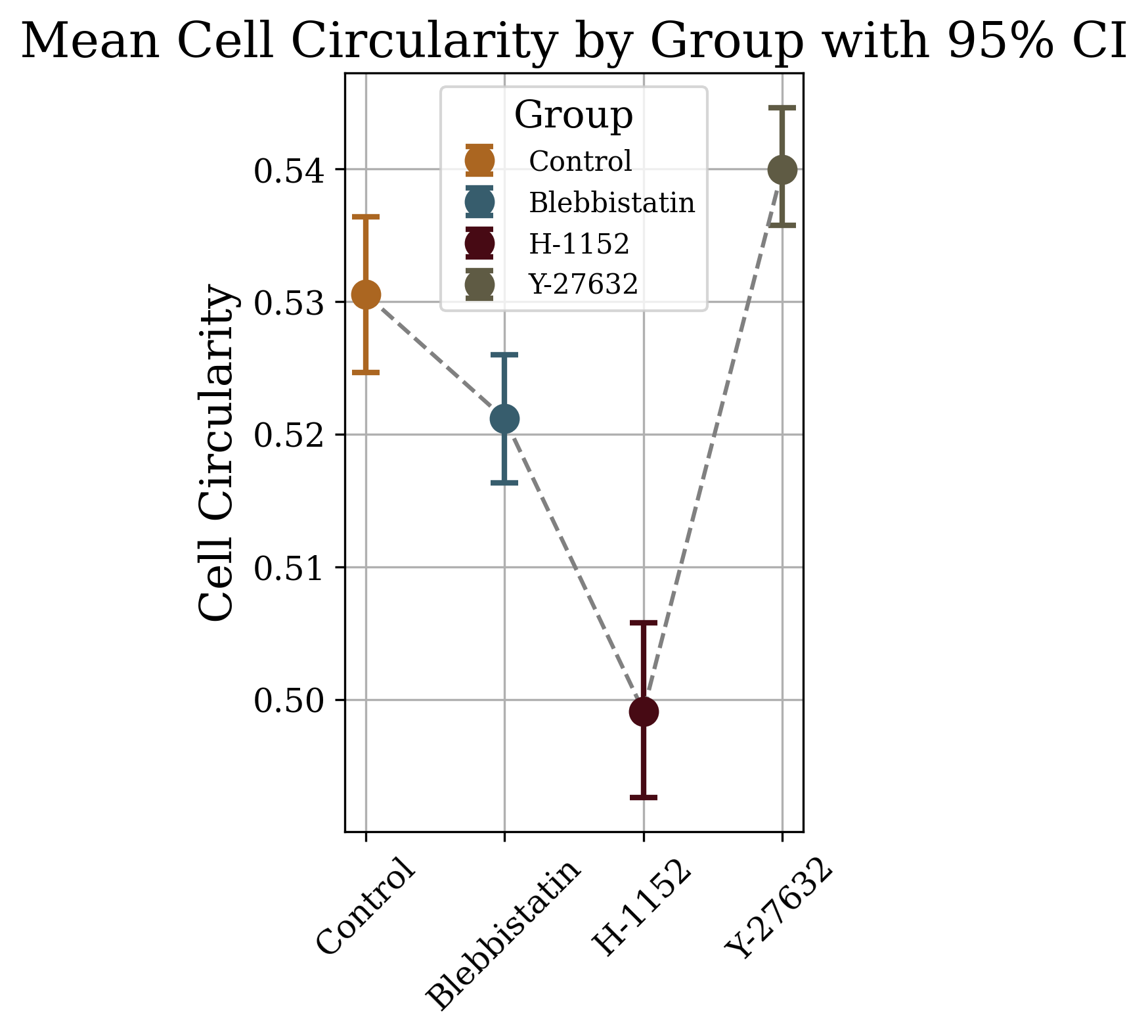

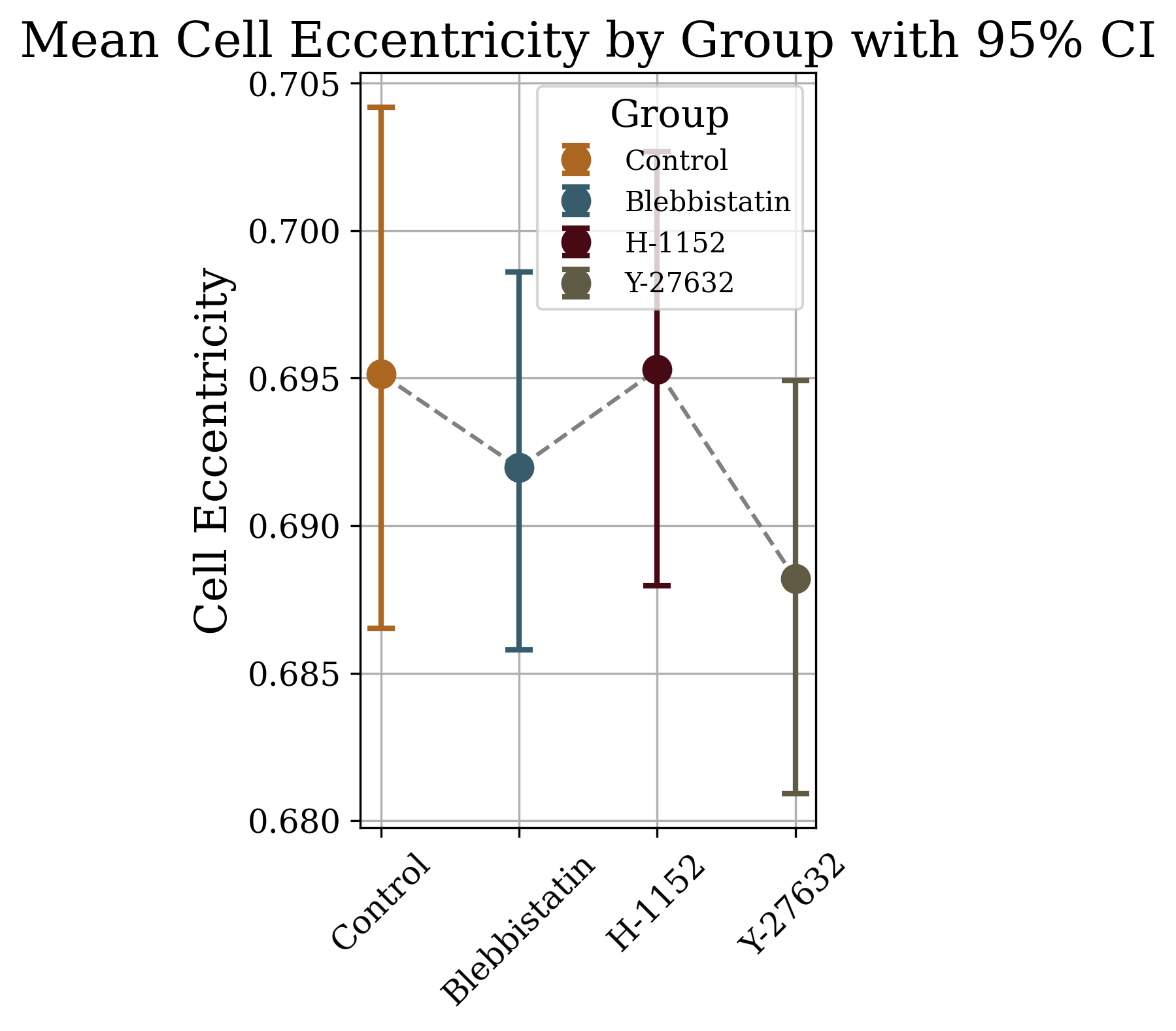

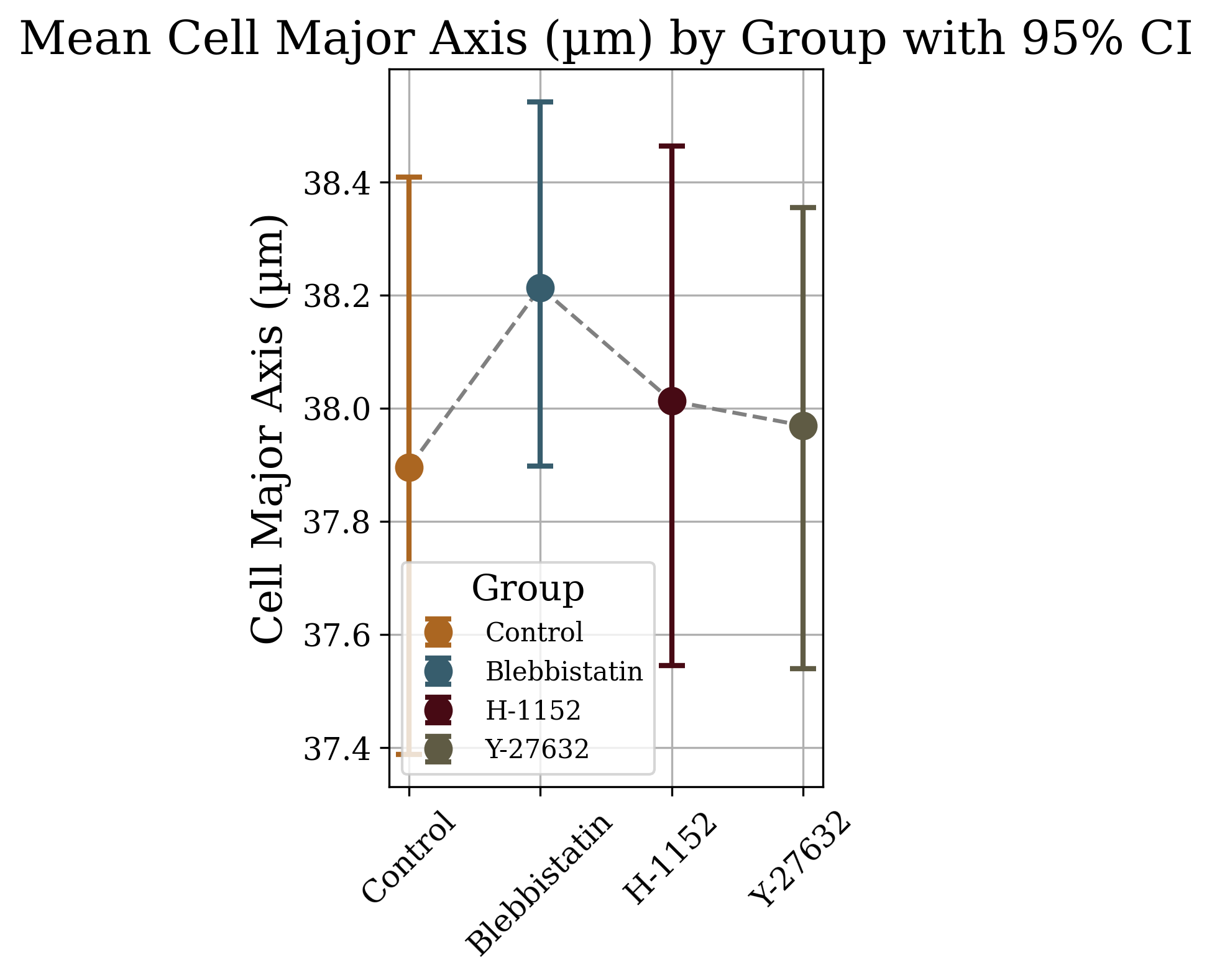

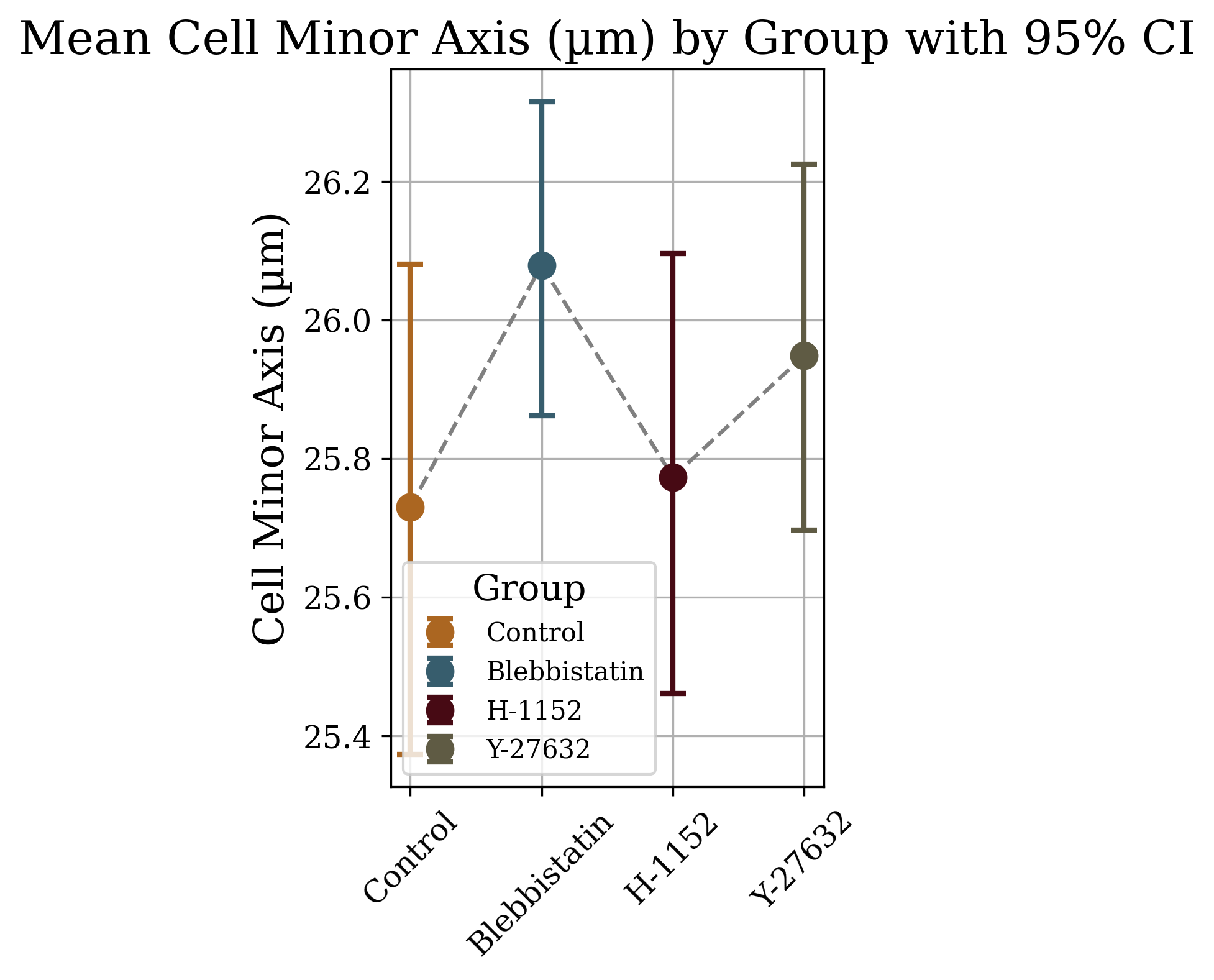

F**igure S3.** Distributions and 95% Confidence Intervals (CI) for select cell features of HeLa cells by treatment group. Far left column contains density distributions of size-related features (cell area, minor and major axes). Middle left are means and 95% CI for each treatment group for the cell features to the left. The middle right column is density distributions for cell circularity, aspect ratio and eccentricity paired with mean and 95% CI for each in the far-right column.

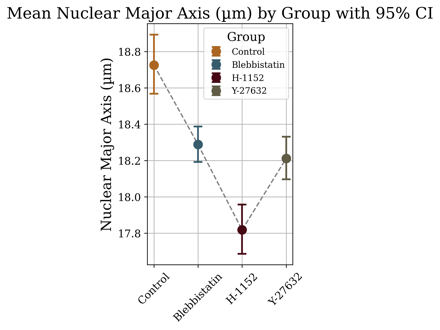

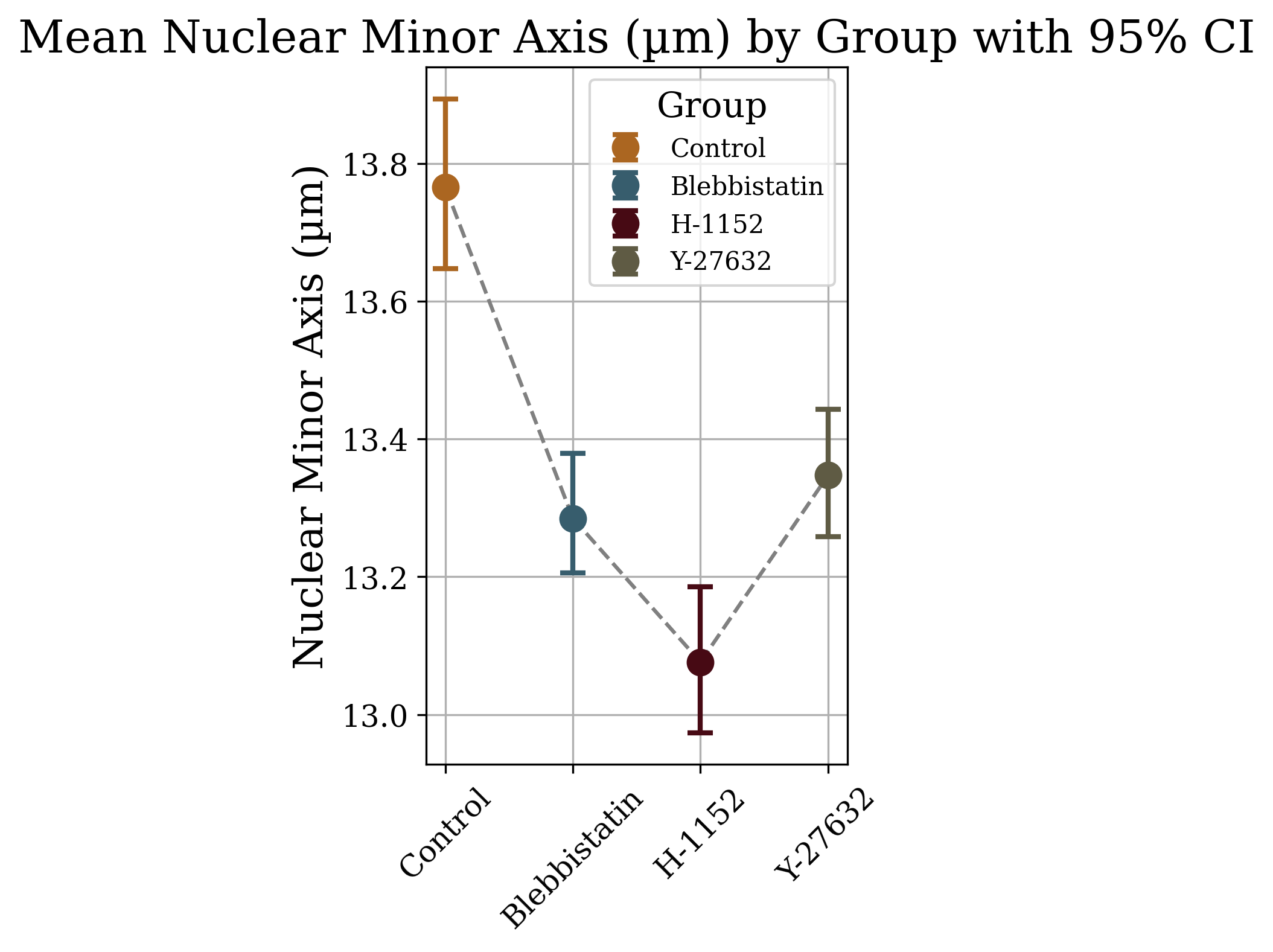

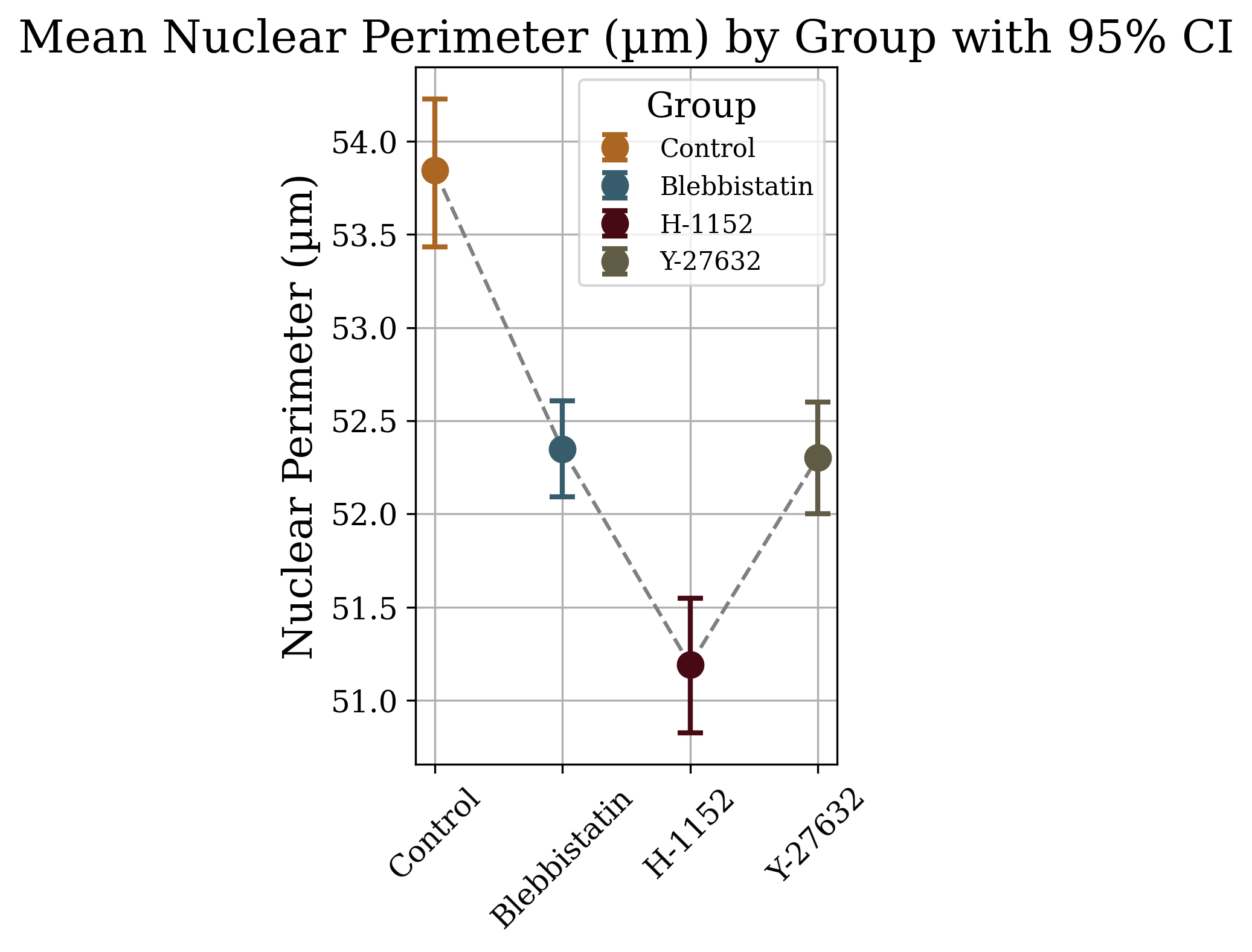

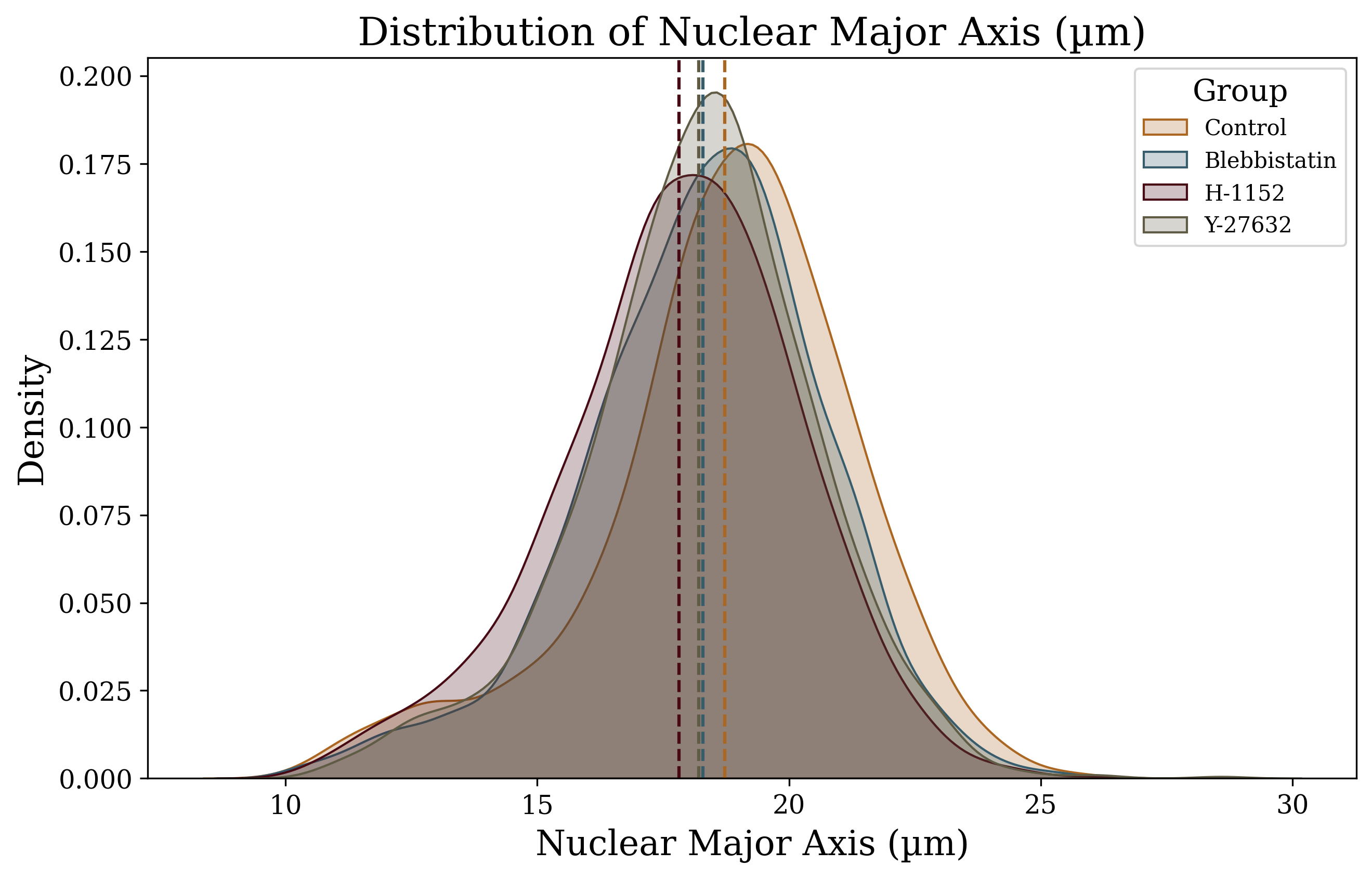

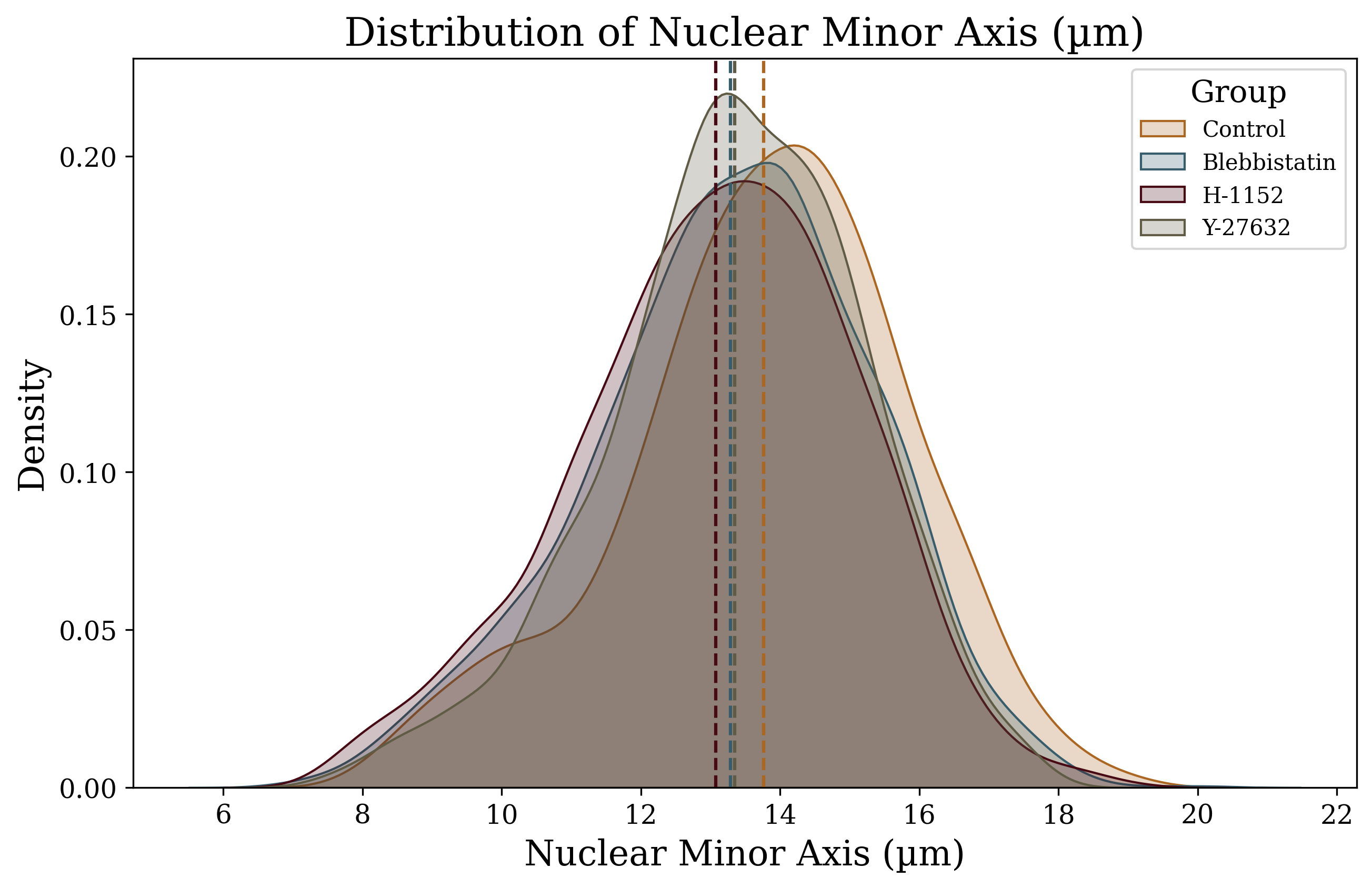

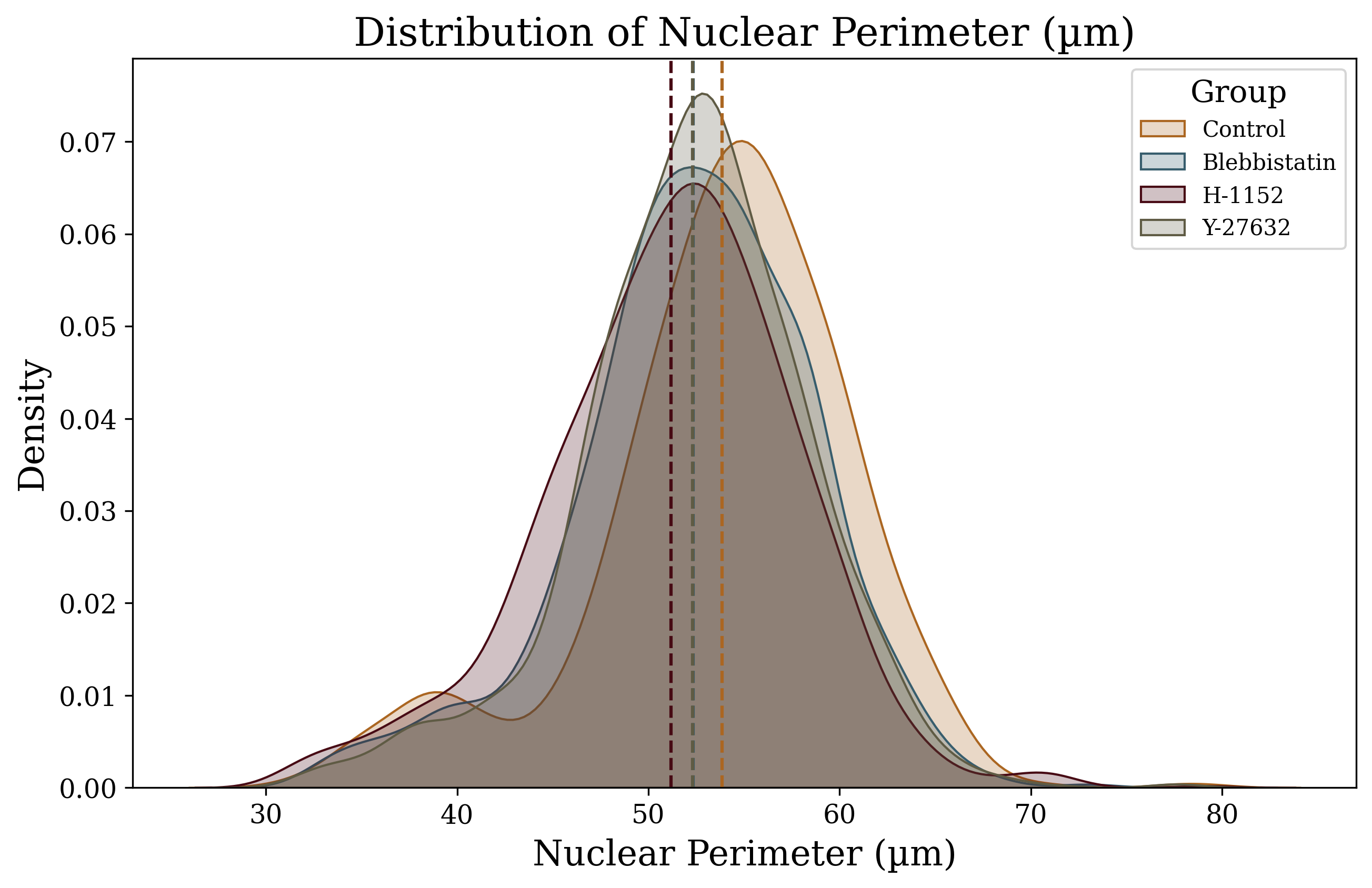

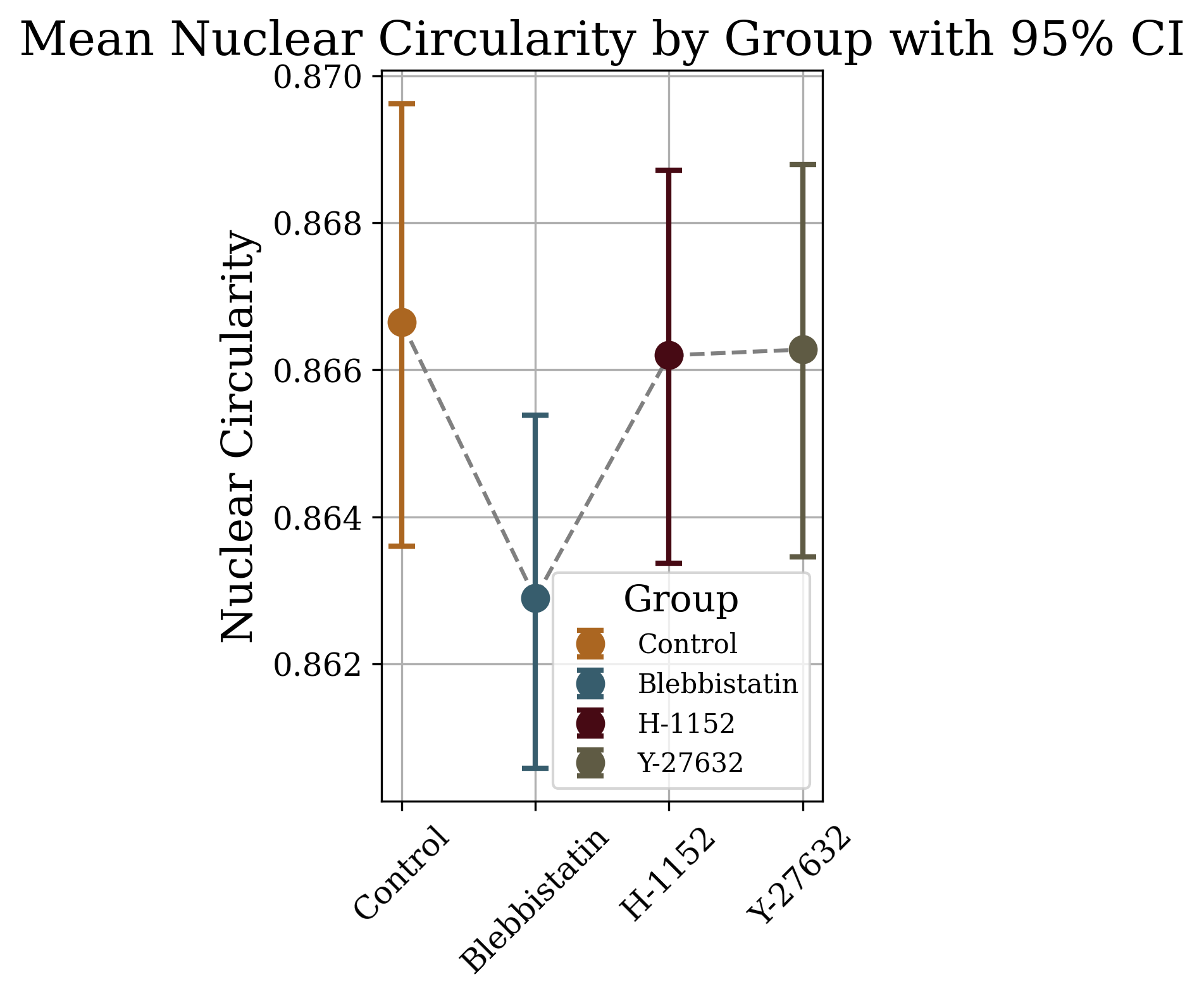

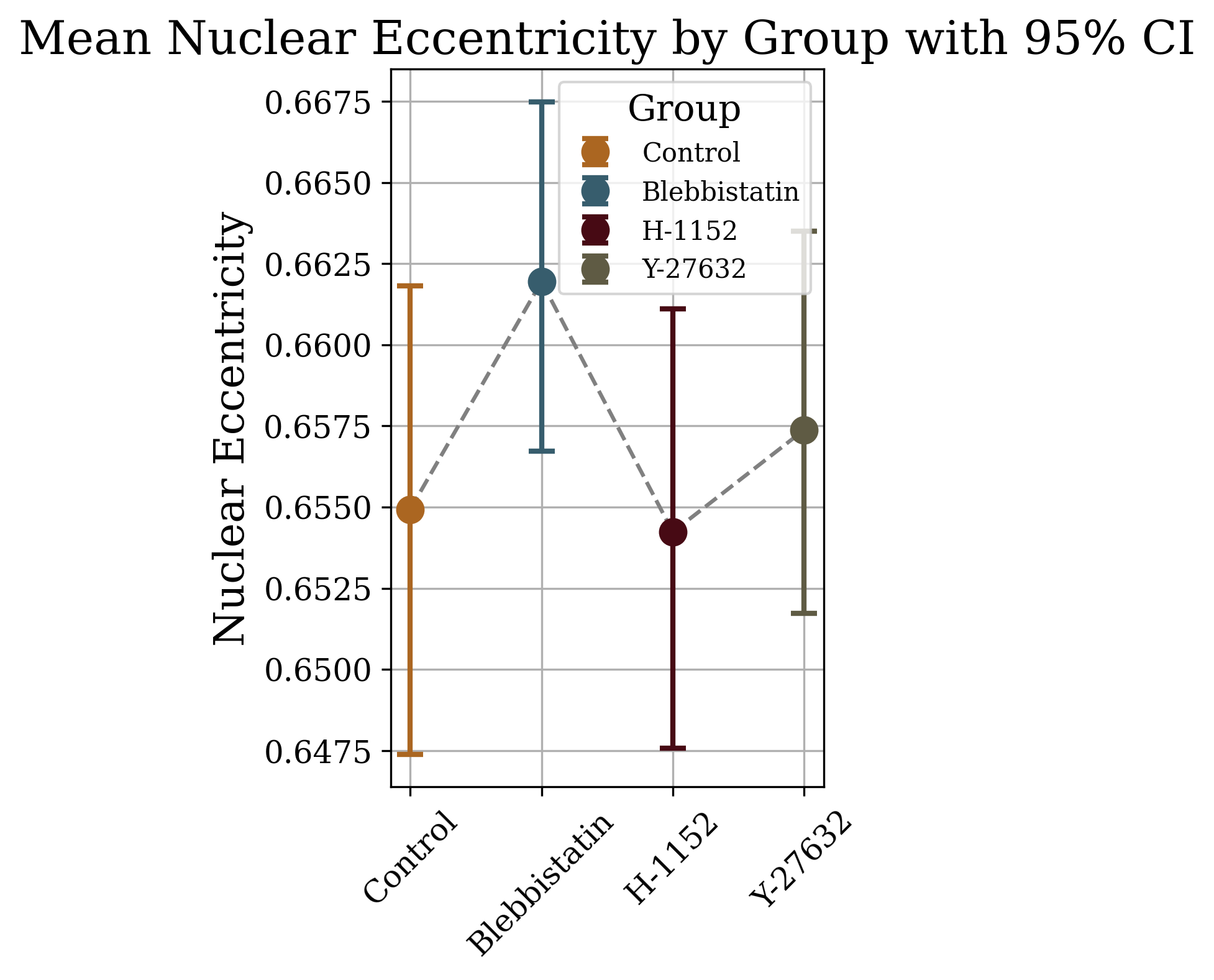

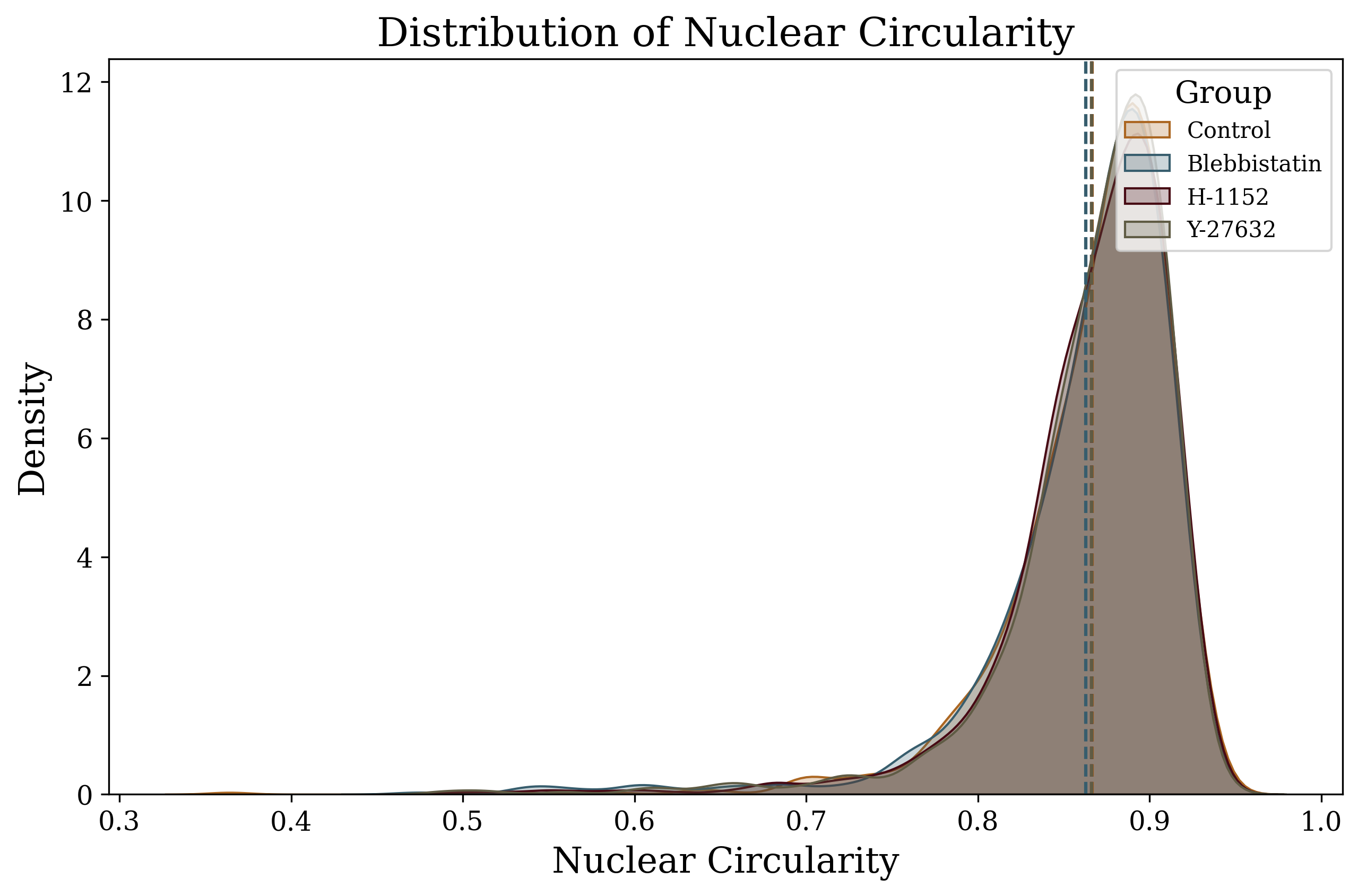

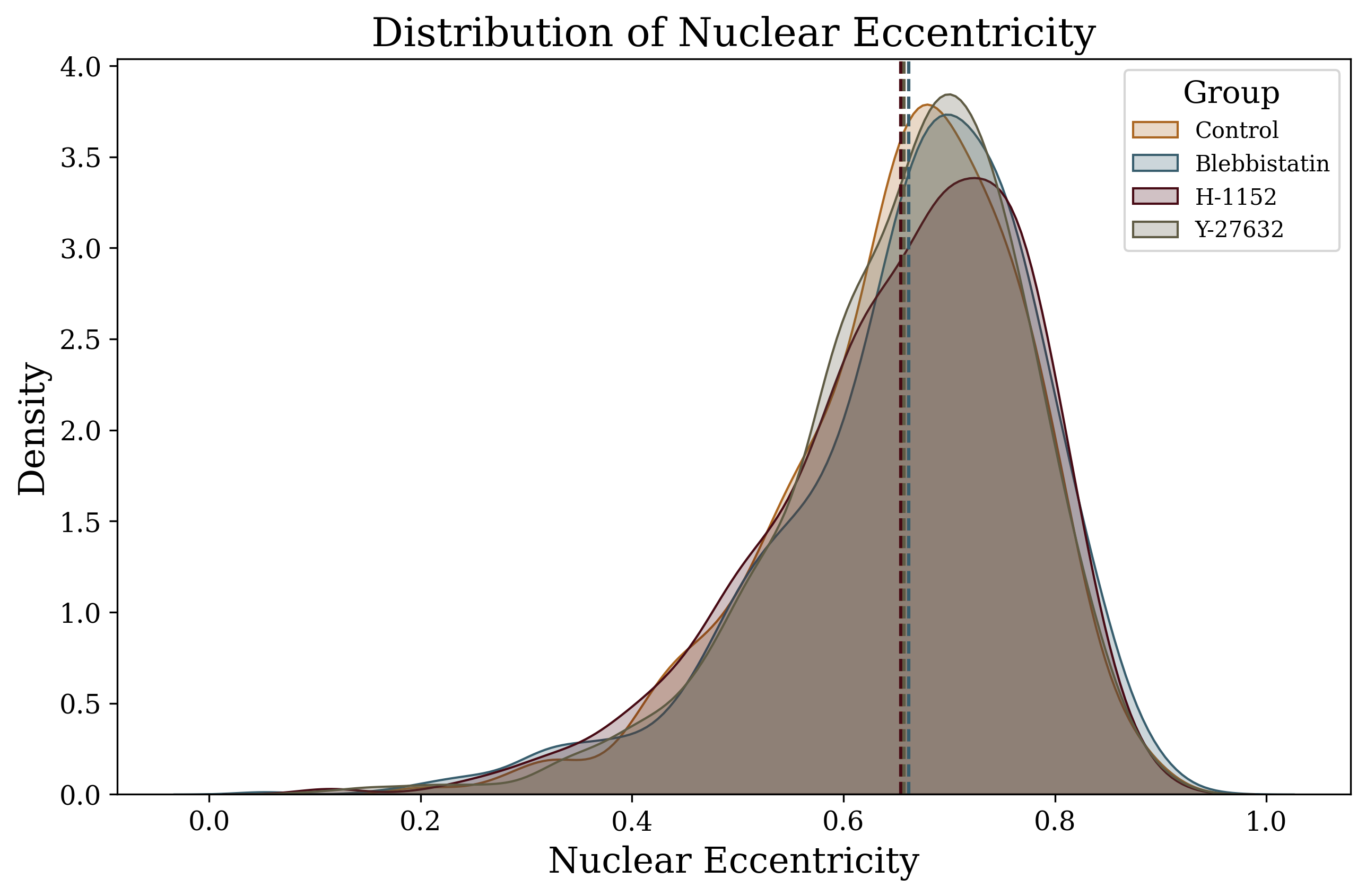

**Figure S4.** Distributions and 95% CI for select nuclear features of HeLa cells by treatment group. Left column are density distributions of size-related features (nuclear minor and major axes, perimeter, circularity, and eccentricity). Right are means and 95% CI for each treatment group.

| **Simple Linear Regression for HeLa cells (Ordinary Least Squares)** | | | | |
| --- | --- | --- | --- | --- |
| **Dependent Feature** | **Group** | **Independent Feature** | **R²** | **p-value** |
| **Nuclear Equivalent Diameter** | |  |  |  |
|  | **Control** | Cell Equivalent Diameter | 0.602566955 | 3.63E-197 |
|  | **Blebbistatin** | Cell Equivalent Diameter | 0.519077412 | 0.00E+00 |
|  | **H-1152** | Cell Equivalent Diameter | 0.620641192 | 7.78E-263 |
|  | **Y-27632** | Cell Equivalent Diameter | 0.560030384 | 7.21E-262 |
|  | **Control** | Cell Mean Radius | 0.595365225 | 2.27E-193 |
|  | **Blebbistatin** | Cell Mean Radius | 0.474765542 | 3.94E-301 |
|  | **H-1152** | Cell Mean Radius | 0.547088867 | 3.62E-215 |
|  | **Y-27632** | Cell Mean Radius | 0.553991433 | 1.48E-257 |
|  | **Control** | Cell Median Radius | 0.593248569 | 2.88E-192 |
|  | **Blebbistatin** | Cell Median Radius | 0.454071599 | 3.45E-283 |
|  | **H-1152** | Cell Median Radius | 0.516221591 | 1.97E-197 |
|  | **Y-27632** | Cell Median Radius | 0.559475479 | 1.80E-261 |
|  | **Control** | Cell Area | 0.534724908 | 7.66E-164 |
|  | **Blebbistatin** | Cell Area | 0.48085999 | 1.50E-306 |
|  | **H-1152** | Cell Area | 0.584938228 | 1.21E-238 |
|  | **Y-27632** | Cell Area | 0.512088313 | 3.78E-229 |
| **Nuclear Area** |  |  |  |  |
|  | **Control** | Cell Equivalent Diameter | 0.589953361 | 1.46E-190 |
|  | **Blebbistatin** | Cell Equivalent Diameter | 0.509901316 | 0.00E+00 |
|  | **H-1152** | Cell Equivalent Diameter | 0.616733916 | 4.44E-260 |
|  | **Y27** | Cell Equivalent Diameter | 0.545514784 | 1.34E-251 |
|  | **Control** | Cell Area | 0.533737818 | 2.15E-163 |
|  | **Blebbistatin** | Cell Area | 0.480010375 | 8.64E-306 |
|  | **H-1152** | Cell Area | 0.591930947 | 3.26E-243 |
|  | **Y-27632** | Cell Area | 0.507455869 | 3.69E-226 |
|  | **Control** | Cell Mean Radius | 0.58436362 | 1.07E-187 |
|  | **Blebbistatin** | Cell Mean Radius | 0.466352764 | 9.41E-294 |
|  | **H-1152** | Cell Mean Radius | 0.546492507 | 8.17E-215 |
|  | **Y-27632** | Cell Mean Radius | 0.54276017 | 1.09E-249 |
|  | **Control** | Cell Median Radius | 0.581123215 | 4.68E-186 |
|  | **Blebbistatin** | Cell Median Radius | 0.445609357 | 4.78E-276 |
|  | **H-1152** | Cell Median Radius | 0.515563882 | 4.58E-197 |
|  | **Y-27632** | Cell Median Radius | 0.548066184 | 2.21E-253 |
|  | **Control** | Cell Maximum Radius | 0.564358971 | 9.29E-178 |
|  | **Blebbistatin** | Cell Maximum Radius | 0.468770977 | 7.33E-296 |
|  | **H-1152** | Cell Maximum Radius | 0.569475937 | 8.35E-229 |
|  | **Y-27632** | Cell Maximum Radius | 0.507395982 | 4.03E-226 |
|  | **Control** | Cell Convex Area | 0.497927329 | 9.59E-148 |
|  | **Blebbistatin** | Cell Convex Area | 0.437876199 | 1.29E-269 |
|  | **H-1152** | Cell Convex Area | 0.553841671 | 3.29E-219 |
|  | **Y-27632** | Cell Convex Area | 0.45677418 | 3.60E-195 |
|  | **Control** | Cell Minor Axis Length | 0.508859634 | 2.12E-152 |
|  | **Blebbistatin** | Cell Minor Axis Length | 0.44143203 | 1.46E-272 |
|  | **H-1152** | Cell Minor Axis Length | 0.536253266 | 8.29E-209 |
|  | **Y-27632** | Cell Minor Axis Length | 0.440545241 | 7.43E-186 |
| **Nuclear Median Radius** |  |  |  |  |
|  | **Control** | Cell Mean Radius | 0.605930595 | 5.79E-199 |
|  | **Blebbistatin** | Cell Mean Radius | 0.48688218 | 5.75E-312 |
|  | **H-1152** | Cell Mean Radius | 0.541075652 | 1.28E-211 |
|  | **Y-27632** | Cell Mean Radius | 0.55421976 | 1.02E-257 |
|  | **Control** | Cell Median Radius | 0.601042478 | 2.34E-196 |
|  | **Blebbistatin** | Cell Median Radius | 0.462106097 | 4.51E-290 |
|  | **H-1152** | Cell Median Radius | 0.505283384 | 2.04E-191 |
|  | **Y-27632** | Cell Median Radius | 0.554838461 | 3.71E-258 |
|  | **Control** | Cell Equivalent Diameter | 0.58399778 | 1.64E-187 |
|  | **Blebbistatin** | Cell Equivalent Diameter | 0.4961016 | 2.19E-320 |
|  | **H-1152** | Cell Equivalent Diameter | 0.588367996 | 7.11E-241 |
|  | **Y-27632** | Cell Equivalent Diameter | 0.520427177 | 1.33E-234 |
|  | **Control** | Cell Maximum Radius | 0.58567264 | 2.30E-188 |
|  | **Blebbistatin** | Cell Maximum Radius | 0.49433994 | 9.17E-319 |
|  | **H-1152** | Cell Maximum Radius | 0.570943358 | 1.01E-229 |
|  | **Y-27632** | Cell Maximum Radius | 0.525199156 | 9.12E-238 |
|  | **Control** | Cell Area | 0.521218662 | 8.62E-158 |
|  | **Blebbistatin** | Cell Area | 0.461256252 | 2.44E-289 |
|  | **H-1152** | Cell Area | 0.553856601 | 3.22E-219 |
|  | **Y-27632** | Cell Area | 0.476931194 | 3.90E-207 |
|  | **Control** | Cell Minor Axis Length | 0.51828754 | 1.68E-156 |
|  | **Blebbistatin** | Cell Minor Axis Length | 0.454317477 | 2.13E-283 |
|  | **H-1152** | Cell Minor Axis Length | 0.528898986 | 1.42E-204 |
|  | **Y-27632** | Cell Minor Axis Length | 0.447528451 | 7.87E-190 |
| **Nuclear Perimeter** |  |  |  |  |
|  | **Control** | Cell Equivalent Diameter | 0.534508692 | 9.60E-164 |
|  | **Blebbistatin** | Cell Equivalent Diameter | 0.45928569 | 1.21E-287 |
|  | **H-1152** | Cell Equivalent Diameter | 0.56081185 | 1.91E-223 |
|  | **Y-27632** | Cell Equivalent Diameter | 0.497817215 | 5.00E-220 |
|  | **Control** | Cell Area | 0.470817076 | 1.27E-136 |
|  | **Blebbistatin** | Cell Area | 0.424588696 | 9.11E-259 |
|  | **H-1152** | Cell Area | 0.529415943 | 7.17E-205 |
|  | **Y-27632** | Cell Area | 0.455130219 | 3.25E-194 |
|  | **Control** | Cell Median Radius | 0.501130533 | 4.25E-149 |
|  | **Blebbistatin** | Cell Median Radius | 0.370635951 | 3.79E-217 |
|  | **H-1152** | Cell Median Radius | 0.451920599 | 7.44E-164 |
|  | **Y-27632** | Cell Median Radius | 0.463233853 | 5.90E-199 |
|  | **Control** | Cell Mean Radius | 0.500699227 | 6.47E-149 |
|  | **Blebbistatin** | Cell Mean Radius | 0.384087618 | 3.51E-227 |
|  | **H-1152** | Cell Mean Radius | 0.474349361 | 4.25E-175 |
|  | **Y-27632** | Cell Mean Radius | 0.454591716 | 6.68E-194 |
|  | **Control** | Cell Maximum Radius | 0.483581396 | 8.72E-142 |
|  | **Blebbistatin** | Cell Maximum Radius | 0.382464987 | 5.85E-226 |
|  | **H-1152** | Cell Maximum Radius | 0.488150681 | 2.95E-182 |
|  | **Y-27632** | Cell Maximum Radius | 0.420468564 | 1.07E-174 |
|  | **Control** | Cell Minor Axis Length | 0.45662981 | 5.03E-131 |
|  | **Blebbistatin** | Cell Minor Axis Length | 0.373331748 | 3.85E-219 |
|  | **H-1152** | Cell Minor Axis Length | 0.470745455 | 2.93E-173 |
|  | **Y-27632** | Cell Minor Axis Length | 0.380430622 | 1.48E-153 |

**Table S3.** Results from linear regression of nuclear shape parameters as a function of cell shape parameters. Table reports R^2^ and p-values for select nuclear-cell feature pairs. Each row is from a linear regression with a single explanatory cell feature variable, as indicated.

| Conover's Test P-values (HeLa) | | | | | | | |
| --- | --- | --- | --- | --- | --- | --- | --- |
|  | **Control** | | **Blebbistatin** | **H-1152** | | | **Y-27632** |
| **Control** | 1 | | 8.94x10^-17^ | 0.000137713 | | | 0.008502463 |
| **Blebbistatin** | 8.94x10^-17^ | | 1 | 7.83x10^-08^ | | | 2.24x10^-14^ |
| **H-1152** | 0.000137713 | | 7.83x10^-8^ | 1 | | | 0.063870371 |
| **Y-27632** | 0.008502463 | | 2.24x10^-14^ | 0.063870371 | | | 1 |
| Conover's Test P-values (NIH3T3) | | | | | | | |
|  | **Control** | **Blebbistatin** | | | **H-1152** | **Y-27632** | |
| **Control** | 1 | 0.72 | | | 5.16 x10^-4^ | 5.87 x10^-7^ | |
| **Blebbistatin** | 7.21 x10^-1^ | 1 | | | 5.99 x10^-4^ | 1.73 x10^-8^ | |
| **H-1152** | 5.16 x10^-4^ | 5.99E-04 | | | 1 | 2.83 x10^-22^ | |
| **Y-27632** | 5.87 x10^-7^ | 1.73E-08 | | | 2.83 x10^-22^ | 1 | |

**Table S4.** Table of Conover’s non-parametric test p-values for the Chromatin Condensation Parameter (CCP) values of each treatment group compared with CCP for all groups.

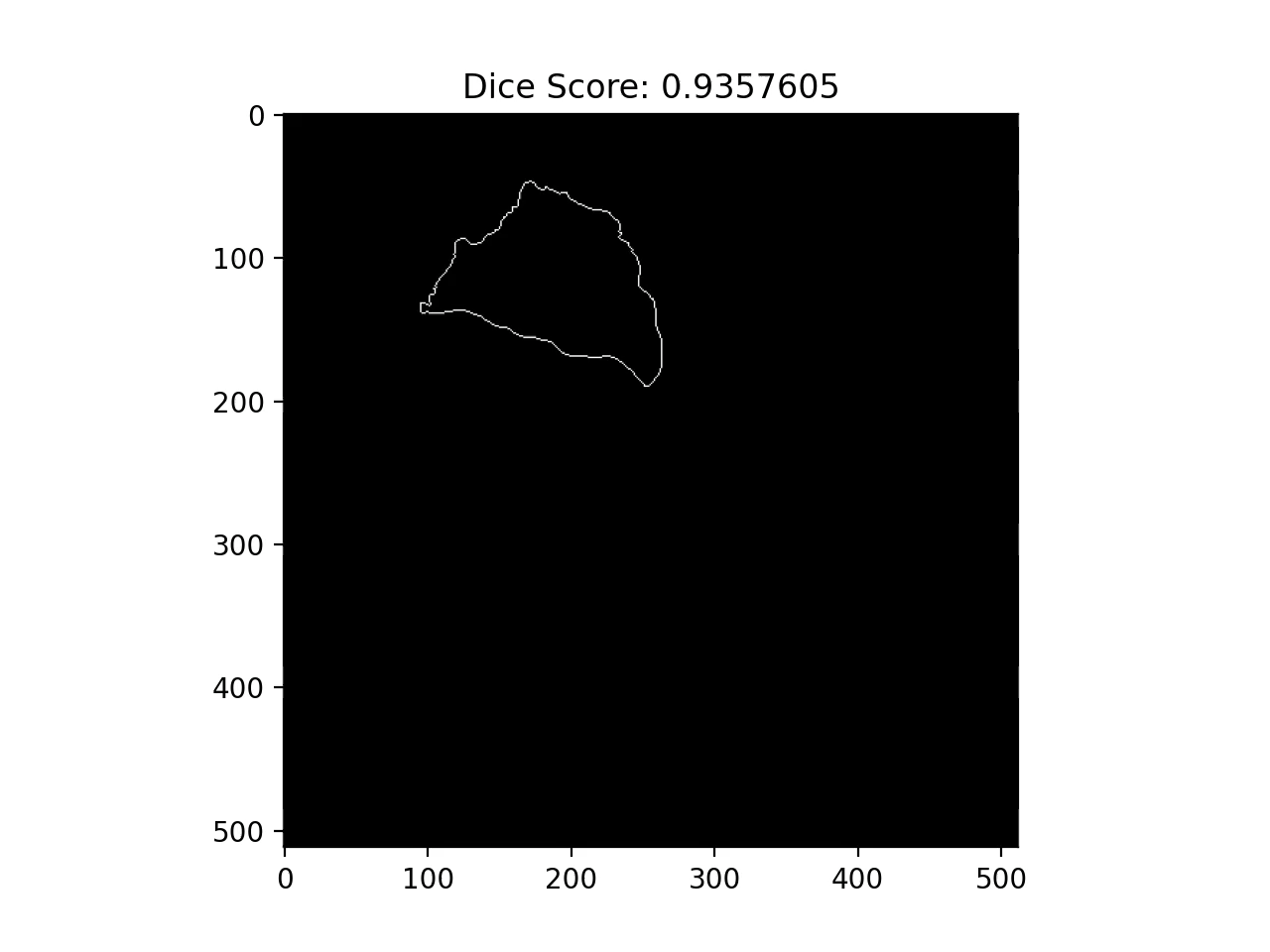

**Figure S5.** An animation that shows a good prediction of the Cell2Nuc model on a hiPSC cell from the Allen Institute image database. This animation shows all the z-stacks of the image, starting from the top of the cell. The pixels forming the cell membrane that were used to generate the prediction form the white boundary. The inner solid pixels represent the nucleus, actual or predicted and the colors have the following meaning. Magenta pertains to nuclear pixels correctly predicted by the model, white pertains to pixels that were not predicted as nucleus but were actually part of the nucleus (false negatives) and green pertains to those pixels that were falsely predicted as belonging to the nucleus (false positives).

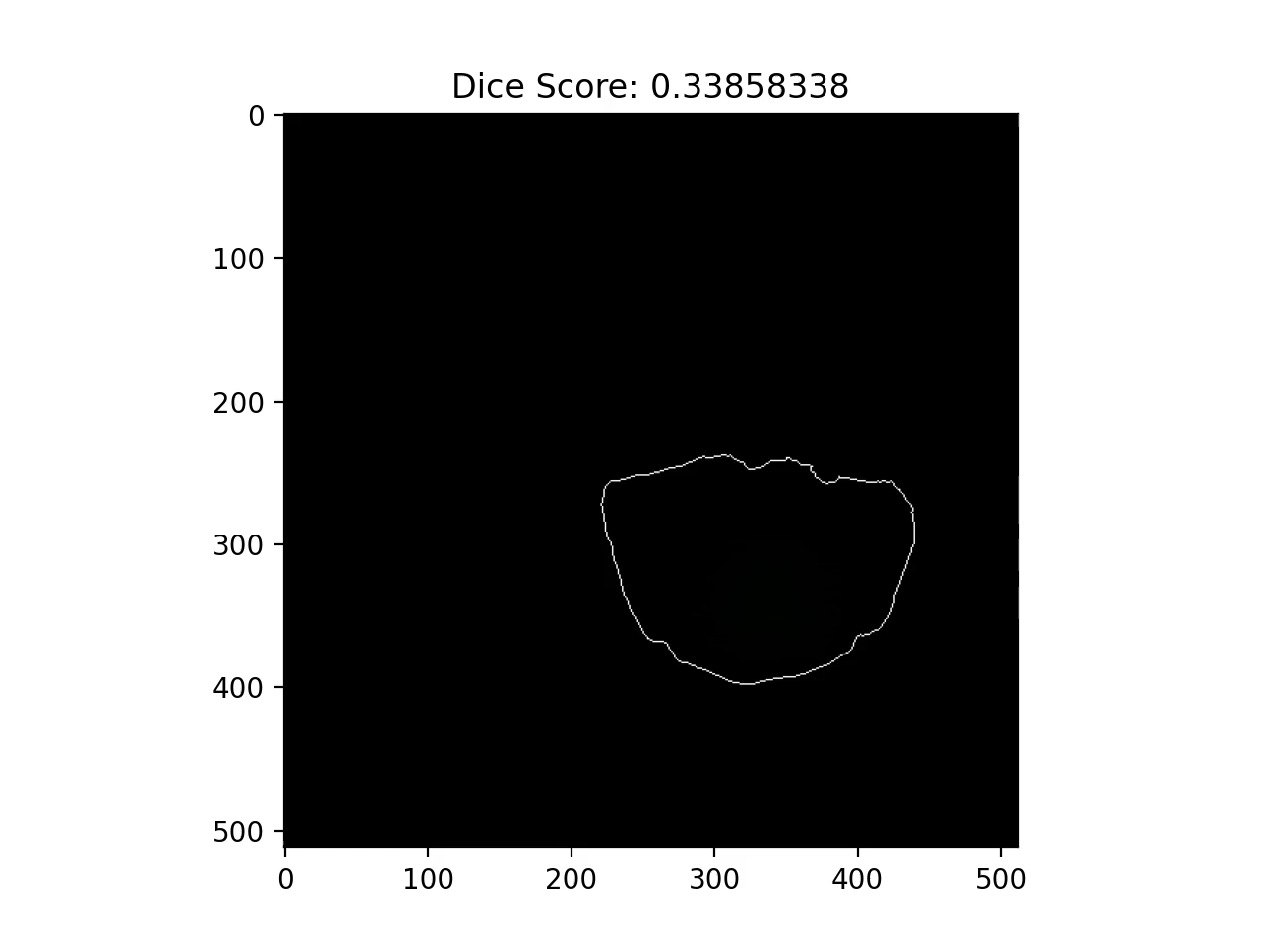

**Figure S6.** An animation that shows a bad prediction of the Cell2Nuc model on a hiPSC cell from the Allen Institute image database. It can be seen here that the nucleus is positioned at one side of the cell and has a non-standard shape for unknown reasons. This animation shows all the z-stacks of the image, starting from the top of the cell. The pixels forming the cell membrane that were used to generate the prediction form the white boundary. The inner solid pixels represent the nucleus, actual or predicted and the colors have the following meaning. Magenta pertains to nuclear pixels correctly predicted by the model, white pertains to pixels that were not predicted as nucleus but were actually part of the nucleus (false negatives) and green pertains to those pixels that were falsely predicted as belonging to the nucleus (false positives).
